# A dimerization-activated proximity labeling system for direct characterization of cadherin *cis* interactions

**DOI:** 10.1101/2025.11.07.687288

**Authors:** Carolyn Marie Orduno Davis, Samveda Shailendra Pagay, Prashant Khatiwada, Yinghao Wu, Sanjeevi Sivasankar

## Abstract

E-cadherins (Ecads) are essential transmembrane cell-cell adhesion proteins that mediate epithelial tissue formation and maintenance. Robust adhesion requires *cis* dimerization between neighboring Ecads on the cell surface and previous structural and biophysical studies have proposed conflicting models on the role of specific and nonspecific interactions in mediating *cis* dimerization. However, since these studies were carried out with isolated Ecad extracellular regions in cell-free systems, it is unknown if specific and nonspecific *cis* binding modes also occur with transmembrane Ecads in live cells, and if specific and nonspecific *cis* dimers recruit different sets of cytoplasmic proteins to Ecad junctions. To directly address these knowledge gaps, we developed a Dimerization Activated TuboID (DAT) proximity labeling system that reports on different modes of Ecad *cis* binding and their corresponding proteomes. Using DAT, fluorescence measurements, diffusion-reaction simulations, and adhesion assays, we demonstrate that Ecads in live cells form *cis* dimers via both specific and non-specific interactions and that *cis* dimerization does not require either prior *trans* contacts or an intact actin cytoskeleton. However, we show that the loss of specific *cis* interactions results in increased junctional instability and Ecad mobility, which leads to dysregulated peripheral protein interactions without affecting recruitment of core junctional proteins. Our study provides key mechanistic insights on cadherin *cis* interactions and also presents a toolkit that can be used to study a broad range of protein *cis* dimers in live cells.

## Introduction

Classical cadherins are a large family of transmembrane cell-cell adhesion proteins that play essential roles in tissue formation and the maintenance of tissue integrity. Deficiencies in cadherin adhesion have been implicated in a number of cancers and epithelial barrier pathologies <u>(Berx and van Roy 2009; Xie et al. 2022)</u>. Among the most widely studied classical cadherin is E-cadherin (Ecad), a prototypic cadherin in epithelia, which mediates epithelial integrity and also functions as a tumor suppressor <u>(Mendonsa et al. 2018)</u>.

Like other classical cadherins, Ecad interacts in two major conformations: a *trans* conformation that enables interactions between Ecads from opposing cells and a *cis* conformation that facilitates binding between Ecads on the same cell membrane. Adhesion is believed to be initiated through the *trans* dimerization of opposing cadherins, which then undergo lateral *cis* clustering through still poorly understood mechanisms. The assemblies that are formed, template the recruitment of cytoplasmic proteins, which may then initiate downstream signaling events <u>(Jamora and Fuchs 2002)</u>. While the structural and biophysical basis for *trans* interactions between opposing cadherins has been extensively studied <u>(Harrison et al. 2010; Parisini et al. 2007; Häussinger et al. 2004)</u>, the molecular mechanisms by which cadherins interact in *cis* orientation are not yet completely understood <u>(Zhang et al. 2009)</u>.

Previous structural studies in canine Ecad have identified that *cis* binding is mediated by specific interaction between two amino acids: a conserved Valine (V81) residue on the outermost (EC1) domain of one cadherin and a Leucine (L175) on the second (EC2) domain of an adjacent cadherin (Fig. 1A) <u>(Harrison et al. 2011)</u>. However, single molecule fluorescence measurements with Ecad ectodomains on supported lipid bilayers, supported by computer simulations, demonstrate that mutating this specific *cis* binding residues does not eliminate Ecad’s ability to form *cis* dimers. Instead, the Ecad mutants continue to *cis*-dimerize via weaker “nonspecific” interactions <u>(Thompson et al. 2020)</u>. However, it is unclear if these ‘specific’ and ‘nonspecific’ *cis* interactions are merely an artifact of measurements on isolated cadherin ectodomains, or if they also occur on the cell surface, where the transmembrane Ecads associate with a large complement of signaling and adaptor proteins and ultimately link to the actin cytoskeleton.

**Figure 1.**
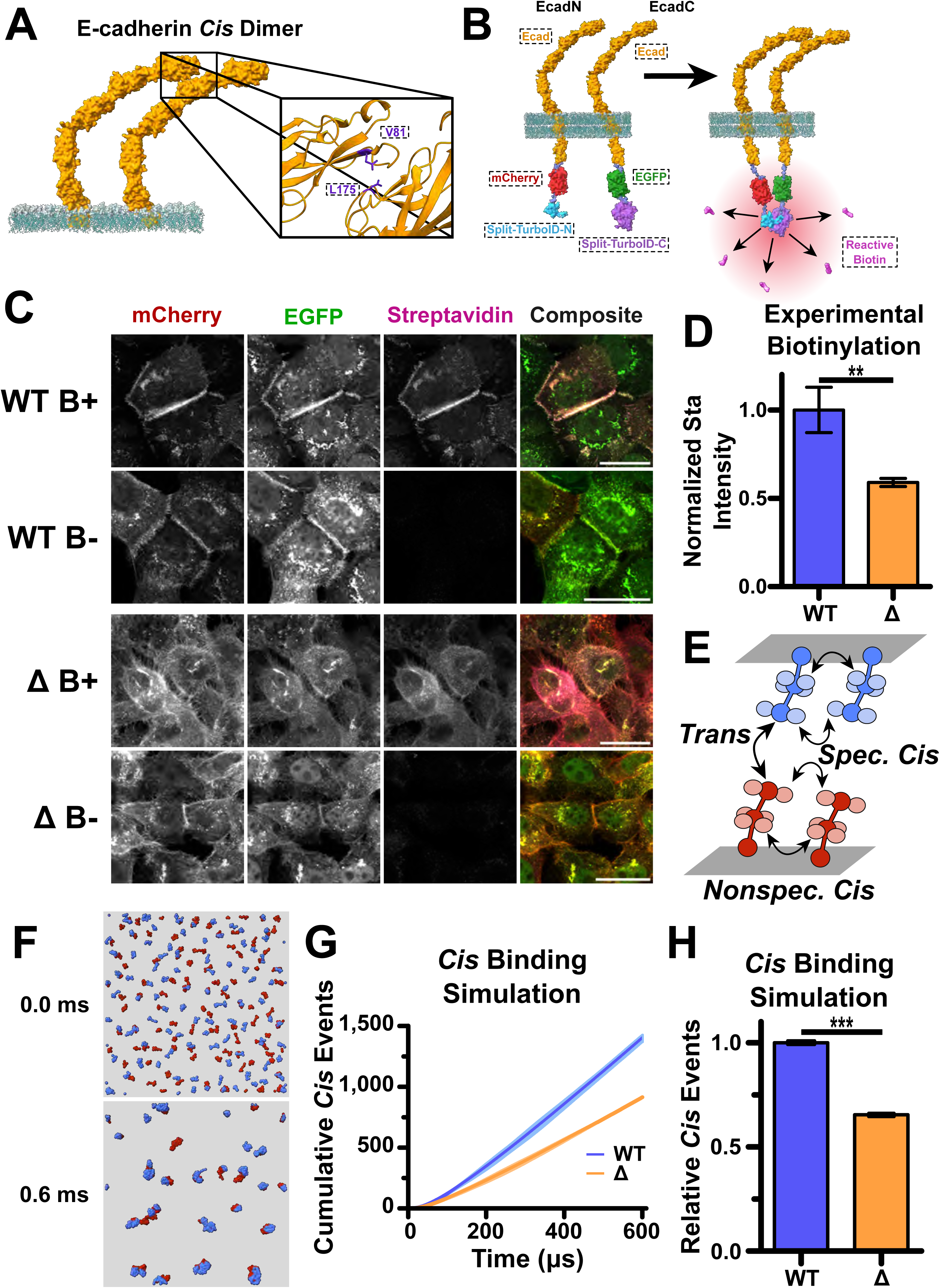
Ecad-DAT provides an accurate readout of *cis* dimerization. **(A)** Schematic of Ecad EC domains in a *cis* binding conformation. Magnification shows closeup of specific *cis* binding site, with relevant residues (V81 on EC1, L175 on EC2) colored indigo. **(B)** Schematic of DAT constructs used in A431(EP)KO cells. mCherry and the N-term piece of Split TurboID were fused to the C-term of one population of Ecad’s extracellular domain using short, flexible linkers (EcadN). GFP and the C-term piece of Split TurboID were fused to the C-term of another population of Ecad using short, flexible linkers (EcadC). When the two populations of Ecad form *cis* dimers, the halves of Split TurboID are brought into registry and reconstituted into a functional enzyme, which biotinylated nearby proteins. **(C)** After incubation with 100 µM biotin for 4 hours, A431(EP)KO cells were fixed and stained for mCherry, GFP, and Streptavidin as proxies for EcadN, EcadC, and biotin respectively. Cells were imaged using three-channel confocal microscopy. Each column in the grid represents a different channel, with the rightmost column showing a composite image of the three channels. Each row represents a different condition, showing cells expressing WT-DAT in the presence of biotin, WT-DAT in the absence of biotin, Δ-DAT in the presence of biotin, or Δ-DAT in the absence of biotin. WT-DAT and Δ-DAT both show strong membrane expression of their components, and strong membrane biotinylation in the biotin(+) conditions with negligible biotinylation in the biotin(-) conditions. **(D)** Graphical comparison of relative biotinylation levels between WT-DAT and Δ-DAT cells, normalized to WT-DAT fluorescence levels to account for differences in DAT expression. Δ-DAT produces 59.0% the amount of biotinylation as WT-DAT when normalized for expression. WT-DAT had n = 413 measured cells distributed over N = 3 biological replicates. Δ-DAT had n = 360 measured cells distributed over N = 3 biological replicates. Error bars show standard deviation. **P<0.01 by two-tailed unpaired-t -test. **(E)** Schematic of Ecad ectodomain *cis* binding simulation. Two populations of Ecad ectodomains, represented by three rigid bodies in a chain (dark spheres), are constrained to two opposite two dimensional planes to represent two cells in contact. *Trans* interactions between opposing colors are allowed via the EC1 analog sphere. Specific *cis* binding sites are represented by two pale spheres on each EC1 sphere, while nonspecific *cis* binding sites are represented by four pale spheres on the middle EC domain. In the WT condition, all binding sites are allowed to make bonds. In Δ condition, specific binding sites on EC1 are blocked so only *trans* and nonspecific *cis* interactions are created. **(F)** Snapshots showing the two populations of Ecad ectodomain models at 0 ms, before binding is allowed to occur, and after 0.6 ms has elapsed, showing Ecad monomers and fully clustered Ecad, respectively. **(G)** Plot of cumulative specific and nonspecific *cis* binding events over time. Curve is average of three rounds of simulation with 95% CI error bars. **(H)** Graph of comparison of total cumulative *cis* binding events for WT and Δ conditions. Δ-Ecad formed 65.4% the amount of *cis* bonds as WT-Ecad. From same data as in G. Error bars show standard deviation. ***P< 0.001 by two-tailed unpaired t-test. Scale bars: 50 µm.

Furthermore, since previous studies on cadherin *cis* interactions have almost exclusively relied on measuring interactions between Ecad ectodomains in a cell-free context <u>(Wu et al. 2010; Harrison et al. 2011; Biswas et al. 2015; Thompson et al. 2020)</u>, it is unclear if specific and nonspecific *cis* interactions have different Ecad proteomes. In other words, it is unclear if the complement of cytoplasmic and transmembrane proteins that interact with Ecad differ when *cis* dimers are formed via specific versus nonspecific interactions. Here, we therefore sought to investigate if the Ecad specific and nonspecific *cis* interactions measured in supported lipid bilayers also occur on the cell surface and determine if these *cis* interactions alter the Ecad proteome.

Many approaches have been developed to map proteomes; however, proximity labeling (PL) techniques are among the most widely used since they are able to detect transient, dynamic protein-protein interactions in the endogenous cellular environment <u>(Qin et al. 2021)</u>. PL is most often performed by genetically fusing a promiscuous biotin ligase enzyme to a protein of interest (the bait). The enzyme generates reactive biotin handles that covalently tag neighboring proteins (the prey) for subsequent detection <u>(Roux et al. 2012)</u>. However, the current generation of PL methods lack spatial and temporal specificity. TurboID, the current state-of-the-art PL biotin ligase tool, while highly efficient, is also highly promiscuous and biotinylates any protein that is in proximity to the bait from the moment it is synthesized, providing reduced spatial specificity <u>(Branon et al. 2018)</u>. Consequently, a typical TurboID assay results in a lot of false positives that muddy the data. To ameliorate this issue, we decided to use a ‘split’ version of TurboID, that has been split into two, non-functional halves, and that needs to reconstitute for labeling to take place <u>(Cho et al. 2020; Schopp et al. 2017)</u>.

To do this we developed Dimerization Activated TurboID (DAT): a split TurboID PL system that reconstitutes upon bait protein *cis* dimerization and initiates biotinylation. We created a pair of split TurboID constructs with each half of split TurboID fused to a different Ecad, creating two populations of split TurboID-tagged Ecad when co-transfected. In this way Split TurboID will only become active when and where Ecad forms *cis* homodimers. By monitoring the extent of DAT-mediated biotinylation in Ecad mutants capable of forming either specific or nonspecific *cis* dimers, we were able to map these dimerization states in cells. Mass spectrometry of the biotinylated proteins also allowed us to investigate the interactomes for Ecad specific and nonspecific *cis* dimers. We supplemented these DAT experiments with diffusion-reaction simulations to investigate the clustering behavior of Ecad on the cell surface via specific and nonspecific *cis* interactions, as well as their clustering at the cell-cell interface through the combined effects of *trans* and *cis* interactions.

By integrating DAT with fluorescence microscopy, diffusion–reaction simulations, and adhesion assays, we investigated specific and nonspecific *cis*-interactions of Ecads on the cell surface for the first time, gaining insights into their corresponding proteomes and their adhesion. We also cataloged the effect of *trans*-interactions and the actin cytoskeleton on Ecad *cis* dimerization, and demonstrate that in cells, Ecads form *cis* dimers via both specific and non-specific interactions. Additionally, we show that the formation of these *cis* dimers does not require either prior *trans* contacts or an intact actin cytoskeleton. However, specific *cis* interactions are critical for stabilizing cell-cell contacts and mediating strong adhesion. Consequently, while the core cytoplasmic protein complex that associates with Ecad is similar when *cis* dimers are formed via either specific or non-specific contacts, specific *cis* interactions result in a more robust proteome since Ecads lacking specific interactions are more mobile and form unstable junctions. Besides the insights that our study provides on the biophysics of cadherin *cis* interactions, we anticipate that the methods we present will have broad applications in studying a wide array of transmembrane protein *cis* dimers on live cell surfaces.

## Results

### Ecad on the cell surface form both specific and nonspecific cis dimers

We developed a genetically encoded Dimerization Activated TurboID (DAT) by fusing GFP and the C-terminal section of Split TurboID (EcadC) or mCherry and the N-terminal section of Split TurboID (EcadN) to the C-terminus of Ecad. We thus created two populations of complementary Ecad-Split TurboID that can be co-expressed in mammalian cells (Fig. 1B). Each fused protein additionally has a short 2 amino acid linker between the C-term of Ecad and the GFP or mCherry fluorophore in addition to a 13 amino acid linker between the respective fluorophore and Split TurboID component. EcadN had an additional V5 tag on the N-term of the Split TurboID (N) component. In order to test the role of specific *cis* interactions mediated by the V81 and L175 residues <u>(Harrison et al. 2011)</u>, we created a V81D and L175D mutant version of EcadC and EcadN to create ΔEcadC and ΔEcadN (ΔDAT).

We co-expressed EcadC and EcadN in Ecad-knockout and P-cadherin (Pcad)-knockout A431 cells (A431(EP)-KO) <u>(Troyanovsky et al. 2021)</u>, where they properly localized to cell membranes (Fig. 1C). Since Split-TurboID demonstrates significant biotin labeling after 1 hour <u>(Cho et al. 2020)</u>, we incubated our cells with 100 µM exogenous biotin for 4 hours and visualized intracellular biotinylation by incubating the cells with fluorescently labelled streptavidin. Upon exogenous biotin incubation, the cells showed clear biotinylation localized at the cell membrane (Fig. 1C). Additionally, there was little to no visible biotinylation in cells incubated without exogenous biotin, showing that endogenous biotin levels do not result in significant labelling.

We repeated this experiment with the Δ mutants, expressing them in A431(EP)-KO cells and visualizing them with and without exogenous biotin (Fig. 1C). After 4 hours of incubation, the Δ mutant constructs produced approximately 60% of the biotinylation of the WT constructs, after normalizing for construct expression (Fig. 1D). Since the Δ mutants cannot engage in specific *cis* binding, this suggested that Ecad *cis*-dimerization on the cell surface occurs via both specific and nonspecific interactions with specific interactions accounting for ∼40% of the cis binding.

To validate the results of our DAT experiments, we performed diffusion–reaction binding simulations, where the extracellular region of each Ecad was represented as three rigid bodies linked by two flexible linkers, allowing conformational changes to be simulated by perturbing the angles and dihedrals between consecutive rigid bodies (Fig. 1E). Specifically, *trans* interactions were formed between the top rigid body of an Ecad molecule on the lower side of the cell–cell interface and the top rigid body of another Ecad on the upper side when the distance between them was shorter than a predefined cutoff value. To model specific *cis* interactions, we assigned a donor site (top two pale spheres) on the surface of the "EC1” rigid body of each Ecad, which can specifically bind to an EC1 acceptor site on a neighboring Ecad. This configuration enabled two adjacent molecules to form lateral associations through these oriented interfaces. For nonspecific *cis* interaction, we assigned four nonspecific binding sites on the surface of the middle rigid body of each Ecad (pale spheres on central body). Any of these sites on one Ecad could interact with any of the corresponding sites on another Ecad when their distance fell below a predefined cutoff value. Consequently, Ecads on the same cell membrane were only allowed to form specific and nonspecific lateral *cis* interactions (WT) or only nonspecific *cis* interactions (Δ), while opposing Ecads were only allowed to form *trans* bonds with each other. In this way the junctional interface between two cells was simulated.

The initial configuration was generated by randomly distributing molecules on a flat surface that represented the plasma membrane. To model the cell–cell interface, two parallel surfaces were positioned facing each other, with the inter-surface distance set to 24 nm, the typical intercellular spacing observed during cell adhesion <u>(Wu et al. 2010; Boggon et al. 2002)</u>. At time t=0 ms, the Ecads were all unbound and freely distributed on the surfaces (Fig. 1F). As the simulation progressed, the system evolved following a previously developed diffusion-reaction algorithm <u>(Su et al. 2021; Su et al. 2020)</u>. The simulation was then allowed to run for 0.6 ms, with frames rendered every µs. The final frame of the simulation (displayed in Fig. 1F) shows the models forming clusters. The cumulative *cis* binding events over time are shown in Fig. 1G, with the comparison of the final number of *cis* binding events between WT and Δ cells shown in Fig. 1H. In qualitative agreement with the experiments, the Δ simulation showed two thirds as many *cis* interactions as the WT simulation, confirming that DAT is indeed only activated by *cis* binding, and the reduced biotinylation seen in the Δ cells can be attributed to fewer *cis* binding events occurring.

### *Trans* interactions have minimal effect on formation of *cis* interactions

Previous simulations with Ecad ectodomains suggest that *trans* dimers drive *cis* interactions, as they limit the movement of Ecad EC domains, which enables *cis* dimerization to take place <u>(Wu et al. 2011)</u>. However, these results conflict with experiments performed on Ecad ectodomains immobilized on supported lipid bilayers, which show that *cis* interactions occur independent of, and precede *trans* dimer formation <u>(Thompson et al. 2021)</u>. We therefore proceeded to test if intercellular *trans* interactions are required for specific and nonspecific *cis* interactions to occur. To do this, we performed experiments with a low density of cells, where the cells would not form intercellular junctions, ensuring that the only Ecad interactions on a cell surface were *cis* interactions (Fig. 2A). In agreement with the cell-free experiments formation <u>(Thompson et al. 2021)</u>, our data showed that even in the absence of *trans* interactions, Δ cells had a little under half the membrane biotinylation of WT cells, showing that Δ cells formed *cis* interactions less than half as often as WT cells (Fig. 2B,C). This is within the margin of error of the comparative levels measured in confluent cells, showing that WT and Δ Ecad form *cis* interactions similarly in the presence or absence of *trans* interactions.

**Figure 2.**
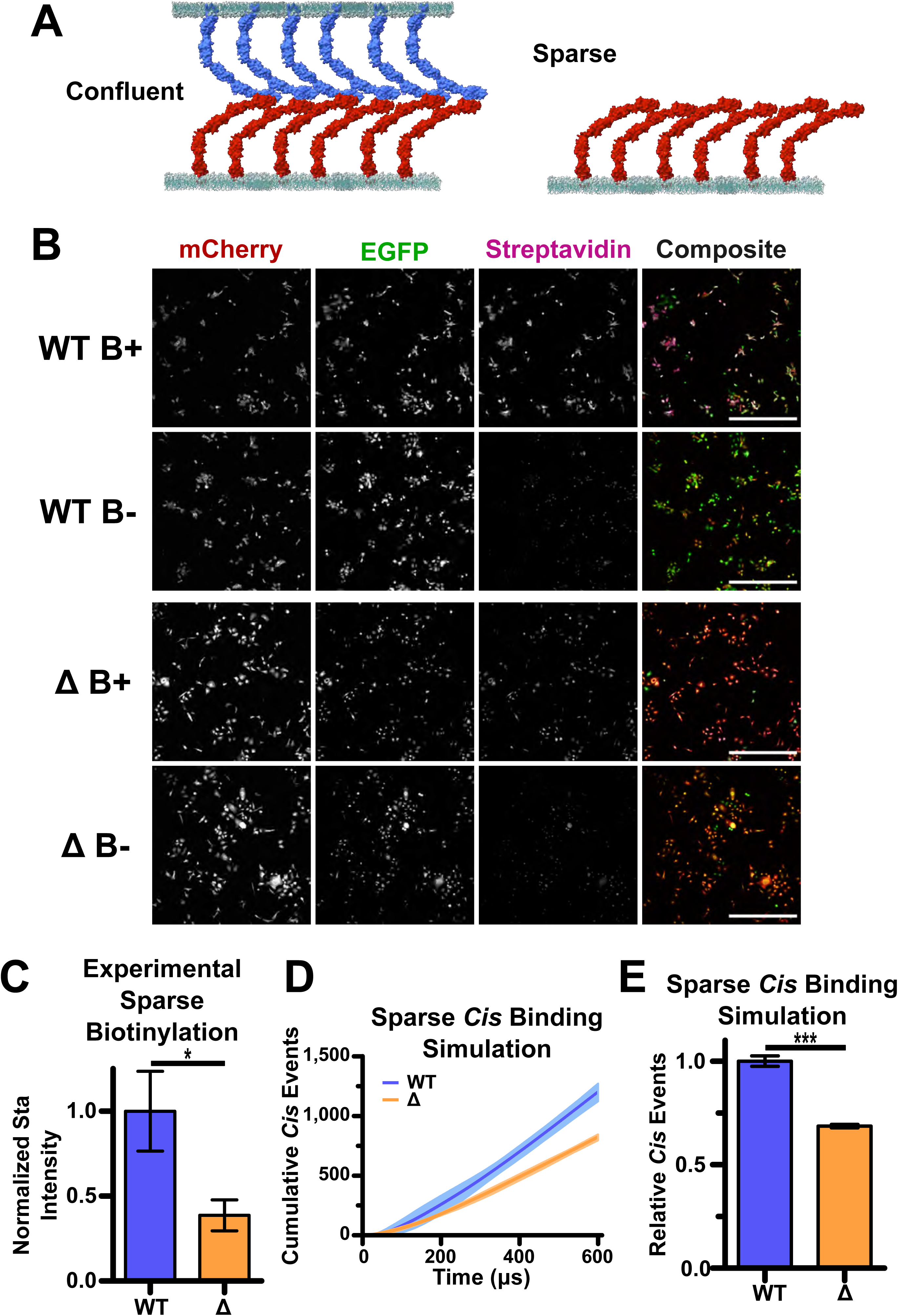
*Trans* Ecad-Ecad interactions affect specific and nonspecific *cis* dimerization equally. **(A)** Schematic comparing the confluent condition, where *trans* interactions can freely occur, with the sparse condition, where cells are physically too far apart for Ecad to form *trans* interactions. **(B)** Cells immunostained for mCherry, GFP, and streptavidin after 4 hours of biotin incubation were imaged with three channel confocal microscopy. Columns correspond to each fluorophore, with the rightmost column being a composite of all three channels. Rows correspond to sparse WT-DAT cells in the biotin(+) and (-) condition and sparse Δ-DAT cells in the biotin(+) and (-) condition. Sparse cells show detectable biotinylation in the biotin(+) condition for both WT-DAT and Δ-DAT cells. **(C)** Graphical comparison of relative biotinylation levels between sparse WT and Δ-DAT cells, normalized to WT-DAT fluorescence levels to account for differences in DAT expression. Δ-DAT produces 38.7% of the amount of biotinylation as WT-DAT when normalized for expression. WT-DAT had n = 96 measured cells distributed over N = 3 biological replicates. Δ-DAT had n = 87 measured cells distributed over N = 3 biological replicates. Error bars show standard deviation. *P< 0.05 by two-tailed unpaired t-test. **(D)** Plot of cumulative specific and nonspecific *cis* binding events over time for the *trans* bond null simulation. Curve is average of three rounds of simulation with 95% CI error bars. **(E)** Graph of comparison of total cumulative *cis* binding events for WT and Δ conditions in the *trans* bond null simulation. Δ-Ecad formed 68.6% the amount of *cis* bonds as WT-Ecad in the absence of *trans* bonds. From same data as in D. Error bars show standard deviation. ***P< 0.001 by two-tailed unpaired t-test. Scale bars: 500 µm.

We also performed a binding simulation similar to that in Figure 1E-H, but with all *trans* interactions blocked. We thus compared the relative formation of specific and nonspecific *cis* interactions in the absence of *trans* bonds. The simulations show a relationship between the WT and Δ conditions that was similar to the “confluent” simulation results, showing that *trans* interactions do not significantly affect the relative incidence of either specific or nonspecific *cis* interactions (Fig. 2D,E).

Together these results imply that, while *trans* and *cis* interactions may have a cooperative effect <u>(Thompson et al. 2021)</u>, Ecad does not require *trans* interactions to form lateral *cis* interactions, and specific and non-specific *cis* interactions are influenced by *trans* interactions to the same degree.

### The actin cytoskeleton does not influence specific and nonspecific *cis* binding

Next, we tested if the presence of the actin cytoskeleton affects specific and nonspecific *cis* interactions. To do this, we used latrunculin to depolymerize actin and measured DAT mediated biotinylation in confluent cells. We used a range of latrunculin concentrations from 0.1-0.2 µg/mL, since actin became significantly depolymerized at latrunculin concentrations > 0.1 µg/mL and was almost entirely eliminated at 0.2 µg/mL latrunculin (Fig. 3A,B; S1A-C). We used phalloidin to visualize actin depolymerization at increasing latrunculin concentrations in both WT and Δ expressing cells (Fig. 3A,B; S1B,C).

**Figure 3.**
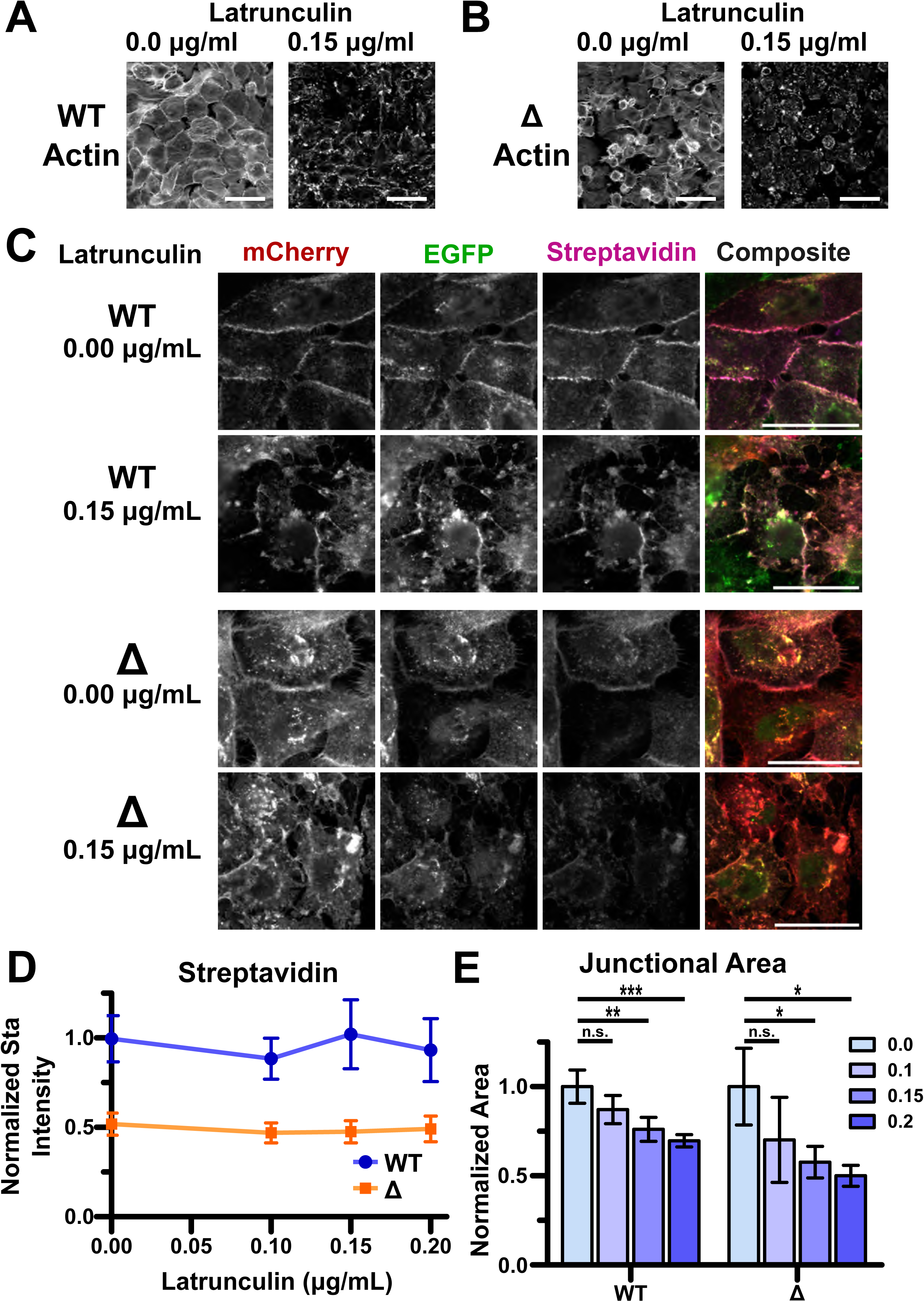
The actin cytoskeleton affects junction size but not Ecad junctional density or *cis* interactions. **(A)** A431(EP)KO cells stabilized with WT-Turbo, stained with phalloidin to detect filamentous actin and imaged in the absence of latrunculin or after half an hour incubation with 0.15 µg/ml latrunculin. After incubation actin structures throughout the cells have nearly entirely disappeared. **(B)** A431(EP)KO cells stabilized with Δ-Turbo incubated in the presence or absence of latrunculin, and stained for actin as in A. After incubation with 0.15 µg/ml latrunculin for half an hour filamentous actin has likewise nearly completely disappeared. **(C)** A431(EP)KO cells expressing WT-DAT or Δ-DAT immunostained for mCherry, GFP, and streptavidin after 4 hours of biotin incubation with and without latrunculin were imaged with three channel confocal microscopy. Columns correspond to each fluorophore, with the rightmost column being a composite of all three channels. Rows correspond to WT-DAT cells in the 0.0 µg/ml and 0.15 µg/ml latrunculin conditions and Δ-DAT cells in the 0.0 µg/ml and 0.15 µg/ml latrunculin conditions. All cells incubated with biotin. All cells show detectable biotinylation in the presence and absence of latrunculin for both WT-DAT and Δ-DAT cells. **(D)** Graph showing streptavidin intensity with increasing latrunculin concentration normalized to Ecad expression levels. Relative to Ecad intensity, streptavidin intensity does not change with increasing latrunculin. n of cells measured are as follows: WT-DAT: 0.0 µg/ml = 260, 0.1 µg/ml = 294, 0.15 µg/ml = 249, 0.2 µg/ml = 249; Δ-DAT: 0.0 µg/ml = 216, 0.1 µg/ml = 205, 0.15 µg/ml = 189, 0.2 µg/ml = 224, distributed over N = 3 biological replicates for all conditions. Error bars show standard deviation. **(E)** Graph comparing relative junctional area with increasing latrunculin concentration. WT-DAT and Δ-DAT junctional areas are normalized to junctional area in the latrunculin negative condition for their respective cell lines. WT junctional area reduced to 69.6% and Δ junctional area reduced to 50.0% under incubation in 0.2 µg/ml latrunculin compared to the latrunculin negative condition. n and N are same as in (d). Error bars show standard deviation. *P< 0.05, **P<0.01, and ***P< 0.001 by two-tailed unpaired t-test. Scale bars: 50 µm.

We also measured the streptavidin intensity at cell junctions at different latrunculin concentrations, and found that there was no change in biotinylation relative to Ecad intensity as latrunculin increased for both WT and Δ cells (Fig. 3C,D; S2A,B). These data suggest that the actin cytoskeleton does not play a significant role in the relative occurrence of specific or nonspecific *cis* interactions.

We also measured the area of cell junctions as latrunculin concentration increased, and found a significant drop as the actin depolymerized (Fig. 3E). Interestingly, the Ecad intensity relative to area did not change, implying that Ecad density remained constant regardless of actin cytoskeleton structure (Fig. S1D), suggesting that the actin cytoskeleton does not affect Ecad density in mature junctions. Interestingly, actin depolymerized at a moderately faster rate in Δ cells (∼40%) compared to WT cells, when normalized for Ecad expression (Fig. S1C). This suggests that specific *cis* interactions may play a role in providing resistance to the effects of latrunculin.

### Specific *cis* interactions result in more robust core junction proteome

Next, we measured the proteome of WT-Ecad and Δ-Ecad using both DAT and full-length TurboID (Turbo). Comparing the proteomes measured using DAT and Turbo enabled us to *(i)* benchmark DAT against Turbo and *(ii)* comprehensively compare the proteomes of WT-Ecad and Δ-Ecad. To measure the Ecad proteome using Turbo, we fused the full-length TurboID enzyme to the C-terminus of either WT-Ecad (WT-Turbo) and Δ-Ecad (Δ-Turbo) and stably expressed these constructs in A431(EP)-KO cells. We incubated WT and Δ cells with 100 μM biotin for 1 hr (Turbo) or 4 hrs (DAT), harvested cell lysate, and performed mass spectrometry on the isolated biotinylated protein.

The proteins found by both WT-Turbo and Δ-Turbo were very similar, with 241 out of 251 proteins detected by WT-Turbo also appearing in the Δ-Turbo results, including major adherens junction proteins known to be in the core Ecad-catenin complex (e.g. α-catenin, β-catenin, p120-catenin, Vinculin, Afadin) (Fig. 4A,B,E) <u>(Guo et al. 2014)</u>. However, Δ-Turbo detected an additional 132 proteins that were not found in the WT results suggesting that Δ-Ecad is more mobile than WT-Ecad and forms less stable junctions. Consequently, Δ-Ecad encounters more proteins which results in a larger proteome (Fig. 4E).

**Figure 4.**
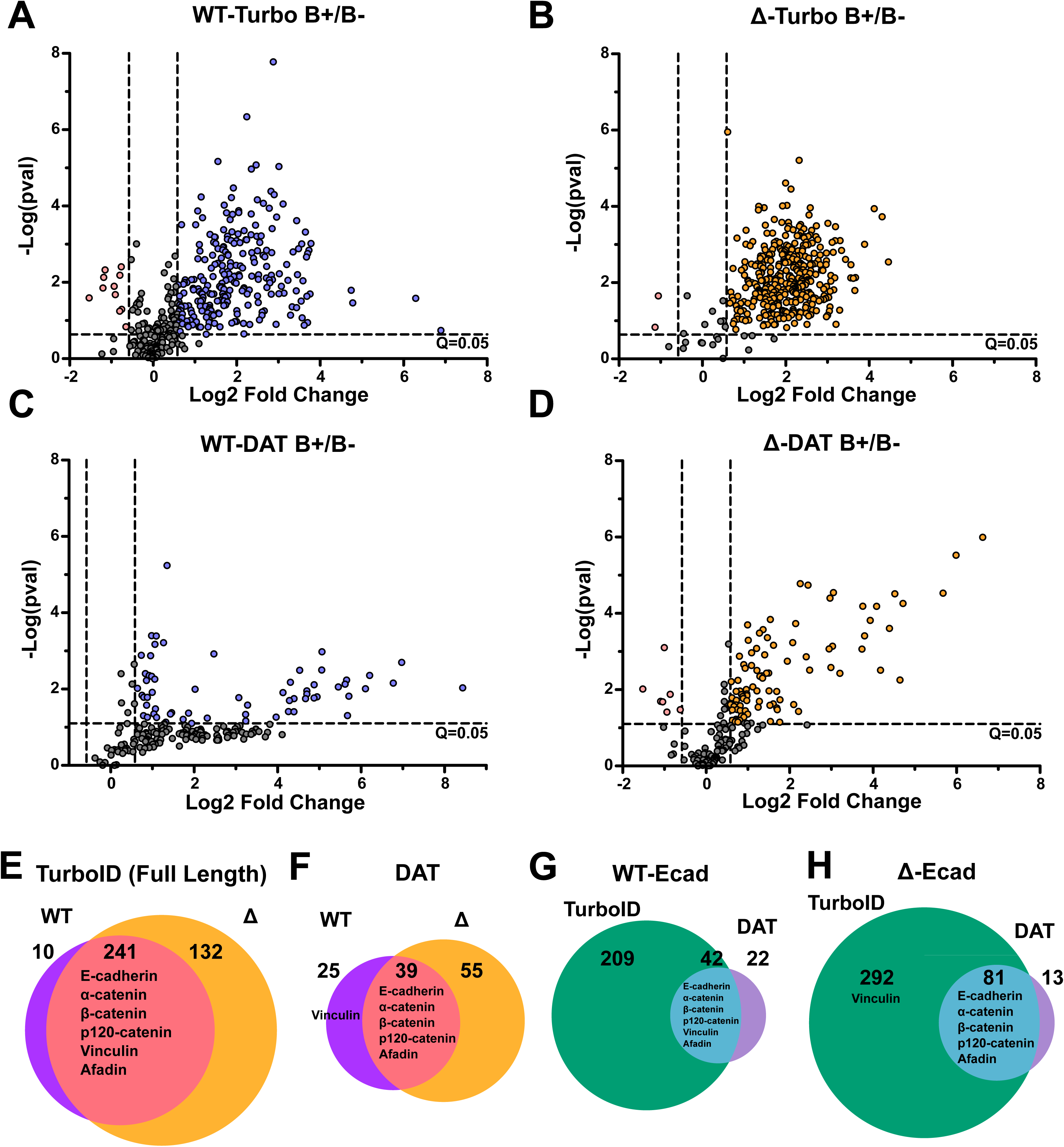
Specific *cis* interactions have little effect on core junctional proteome. Volcano plots showing the enrichment of biotinylated proteins in **(A)** WT-Turbo, **(B)** Δ-Turbo, **(C)** WT-DAT, and **(D)** Δ-DAT in the presence (upper right quadrant) and absence (upper left quadrant) of biotin. Proteins in the upper right quadrant both had a greater than 1.5-fold increase between the biotin(-) and biotin(+) conditions and a greater than 95% confidence of identification. **(E-H)** Venn diagrams comparing the incidences of individual proteins across systems. Δ conditions returned a wider variety of hits than WT cells, and DAT systems returned a smaller set of proteins that nonetheless largely overlapped with the Ecad-Turbo sets. N = 3 biological replicates. Results are compiled in Candidate Lists document.

Cells with both WT-DAT and Δ-DAT likewise had large overlaps in their detected proteomes with 39 out of 64 hits in the WT condition also found in the Δ condition, with both cell populations also showing most of the same important adherens junction proteins, with the exception of vinculin, which was only present in the WT condition (Fig. 4C,D,F). Similar to the Turbo results, Δ-DAT returned 55 unique proteins not present in the WT condition (in contrast to WT-DAT which only found 25 unique proteins), again suggesting that Δ-Ecad has a larger proteome (Fig. 4F).

Comparing the proteomics results for WT-Turbo and WT-DAT, we see that while WT-Turbo returned a broader array of hits, the majority of WT-DAT’s results were shared (Fig. 4G). Of these shared hits, WT-DAT had a near-universally higher fold-change between the biotin (-) control and the biotin (+) experimental condition, showing that WT-DAT makes up for its lower efficiency with lower background labelling and a higher dynamic range (Fig. S3A). This finding was repeated in the comparison between Δ-Turbo and Δ-DAT, though to a lesser degree, with Δ-DAT only having a higher dynamic range on some of the shared hits (Fig. 4H, Fig. S3B). This could again be due to Δ-Ecad having a more variable proteome and/or less stable junctions, creating a less consistent environment for Δ-DAT to label. As anticipated, WT-DAT had consistently higher fold changes than Δ-DAT in their shared hits (Fig. S3C). WT-Turbo and Δ-Turbo had largely comparable signals on a protein-to-protein basis (Fig. S3D).

Additionally, we used the STRING proteomics analysis tool to map out the known interactions in our proteome (Fig. S4) <u>(Szklarczyk et al. 2023)</u>. While WT-DAT returned a tight web of interactions, with only a few major interaction groupings and 12 “isolated” hits with no documented direct interactions (Fig. S4A), Δ-DAT showed a broader web of hits, with small unconnected “pockets” of proteins not generally associated with adherens junctions, in addition to 25 isolated protein hits (Fig. S4B). These results again indicate that a lack of specific *cis* interactions causes improper Ecad localization. Alternatively, the reduction in *cis* interactions could create a less organized junction that allows unrelated proteins access to the normally blocked off area.

We also performed metabolic pathway analysis using the Reactome online tool to analyze the changes in expression levels between WT and Δ cells (Fig. S5, Reactome Pathways) <u>(Milacic et al. 2024)</u>. By wholistically analyzing the proteome changes, Reactome determined that the cell-cell junction organization pathway, particularly the Adherens Junction interaction pathway, was significantly downregulated in the Δ-Ecad cells. Other regulation pathways were also reduced in Δ-Ecad cells. These changes could translate to less organized junctions and weaker cell adhesion.

### Specific *cis* interactions are required for robust Ecad adhesion and clustering

To test the effects of specific *cis* bonds on cell-cell adhesion and clustering, we performed a range of adhesion assays and fluorescence recovery after photobleaching (FRAP) experiments. In order to ensure more consistent Ecad expression in these experiments, we used cells expressing either WT-Turbo or Δ-Turbo constructs.

We first tested the adhesion of cells expressing WT-Ecad or Δ-Ecad using a dispase assay where a confluent monolayer of cells was detached from a substrate using dispase protease and then subjected to mechanical agitation. While cell sheets expressing WT-Turbo remained intact, cells expressing Δ-Turbo fragmented (Fig. 5A-B). We measured a 100-fold higher fragmentation of the Δ-Turbo cell sheets compared to WT-Turbo cell sheets (Fig. 5A-B). Parental A431(EP)-KO cells, which served as a cadherin-negative control, produced 1000-fold more fragments than the WT-Turbo cells (Fig. 5A-B). These experiments showed that specific *cis* interactions are essential in mediating robust cell-cell adhesion, and while nonspecific *cis* interactions are strong enough to form a cohesive cell sheet, the bonds cannot hold up to force nearly to the extent of cells with both specific and nonspecific *cis* interactions.

**Figure 5.**
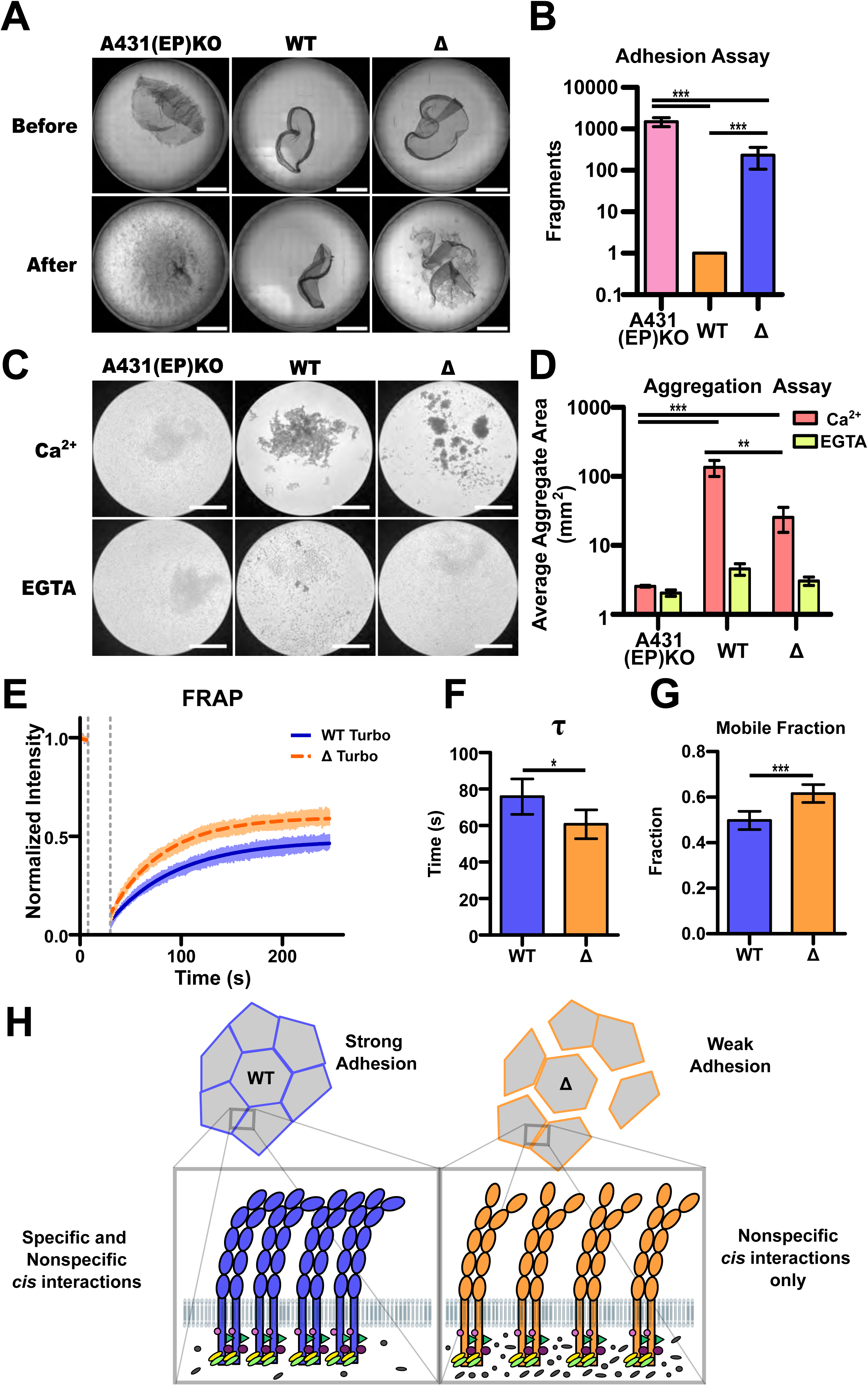
Specific *cis* interactions affect cell adhesion strength, and Ecad mobility. **(A)** Adhesion assay performed using dispase to detach confluent cells into a single monolayer. After applying mechanical stress, the monolayer of A431(EP)KO and Δ-Turbo cells fragmented. WT-Turbo cell monolayer remained intact. Scale bars: 5 mm. **(B)** Quantification of adhesion assay showing number of fragments on a log scale. N = 3 biological replicates. Error bars show SEM. ***P< 0.001 by two-tailed unpaired t-test. **(C)** Cell aggregation assay. Cells were allowed to form aggregates under Ca^2+^ and EGTA. A431(EP)KO cells did not aggregate under either condition, while WT- and Δ-Turbo cells formed aggregates only in Ca^2+^. Scale bars: 2.5 mm. **(D)** Quantification of aggregation assay showing average aggregate area on a log scale. WT-Turbo cells covered nearly an order of magnitude more area than Δ-Turbo cells. N = 3 biological replicates. Error bars show SEM. **P<0.01, ***P< 0.001 by two-tailed unpaired t-test. **(E)** Graph of normalized fluorescence intensity over time. A small segment of WT- or Δ-Turbo cell membrane was bleached for 10 seconds, and fluorescence was allowed to recover until steady state was reached (250 s). Error bars show 95% CI. n = 49 (WT) or 50 (Δ) runs distributed across 3 biological replicates. **(F)** Comparison of the t values. WT cells had a slightly higher t, indicating slower protein migration. Error bars show 95% CI. *P< 0.05 by two-tailed unpaired t-test. **(G)** Comparison of the mobile fraction. WT cells had a smaller mobile fraction than Δ cells, indicating more stable junctions. Error bars show 95% CI. ***P< 0.001 by two-tailed unpaired t-test. **(H)** Schematic of proposed model. Combination of specific and non-specific *cis* interactions are in play for normal Ecad binding. The loss of specific *cis* interactions results in partial junctional instability and increased junctional mobility, leading to a larger interactome without affecting the recruitment of core junctional proteins.

We also tested the ability of WT-Turbo cells and Δ-Turbo cells to form aggregates in the presence of calcium (Ca^2+^ condition) or the presence of EGTA, a Ca^2+^ chelator (EGTA condition). In the presence of calcium, WT-Turbo cells formed large aggregates, with nearly all visible cells contained in an aggregate (Fig. 5C,D). In contrast, Δ-Turbo cells formed smaller aggregates that occupied only half as much area as WT cells (Fig. 5C,D). Parental A431(EP)-KO cells did not form aggregates in the presence of calcium. All cells failed to form aggregates in the calcium-negative EGTA condition (Fig. 5C,D). These experiments again confirm the essential role of specific *cis* interactions in Ecad adhesion.

We additionally used FRAP experiments to investigate the effect of Δ mutations on junctional Ecad mobility. After bleaching a small ∼1 µm segment of the cell-cell junction in WT-Turbo or Δ-Turbo cells, we allowed the fluorescence to recover for 200 s, until it reached steady state (Fig. 5E). The average recovery curves were used to calculate the recovery half-life (t) and the Ecad mobile fraction (Fig. 5F,G). In agreement with previous FRAP experiments on the Δ mutations <u>(Harrison et al. 2011)</u>, we measured a decreased t as well as an increased mobile fraction in Δ cells compared to WT cells. Taken together, our results show that specific *cis* interactions stabilize Ecad at cell-cell junctions and are major contributors to junctional strength. In the absence of these interactions, the Ecad junctions are more mobile and form much weaker contacts.

## Discussion

Cadherin *cis* interactions have been historically difficult to study, since techniques such as atomic force microscopy, which are widely used to measure direct cadherin binding interactions, can only test *trans* interactions. Similarly, fluorescence-based studies of cadherin *cis* interactions and *in silico* simulations have only used the ectodomain, which doesn’t take into account the contributions of the cadherin intracellular region or linkage to the cytoskeleton <u>(Wu et al. 2010; Harrison et al. 2011; Biswas et al. 2015; Thompson et al. 2020)</u>. Consequently, the exact mechanisms of *cis* interactions are still a mystery, although one specific interaction site, the V81 and L175 residues, was experimentally determined to contribute to Ecad *cis* interactions <u>(Harrison et al. 2011)</u>. However, this is not the only mode of *cis* interaction, since mutating these residues do not completely abolish *cis* binding or junction formation <u>(Biswas et al. 2015)</u>. This specific interaction site is nonetheless useful to study, as mutating it was shown to significantly reduce Ecad association and increase its mobility <u>(Harrison et al. 2011)</u>. Therefore, we sought to investigate the contribution of specific and nonspecific interaction to *cis* dimerization in live cells. Here we demonstrate, for the first time, that Ecad *cis* dimerization on the cell surface occurs through both specific and nonspecific *cis*-interactions. We show that specific *cis* interactions are critical for clustering cadherins and mediating strong cell-cell adhesion. Consequently, while the core cytoplasmic protein complex that associates with Ecad is similar when *cis* dimers are formed via either specific and non-specific contacts, the specific *cis* interactions result in a more robust proteome since Ecads lacking specific interactions are more mobile and form unstable junctions (Fig 5H).

In order to investigate Ecad homophilic lateral interactions in more endogenous conditions, we created a tool using the complete transmembrane Ecad, not just the ectodomain, and using split-TurboID to both study the binding itself as well as the proteome that results due to Ecad *cis* dimerization. To this end we created DAT by fusing split-TurboID to two populations of Ecad to get a both direct readout of Ecad-Ecad dimerization via biotinylation, and to obtain a labeled proteome for identification via mass spectrometry. DAT allowed us to determine that removing previously characterized “specific” *cis* interactions reduces Ecad biotinylation (i.e. *cis* binding) by approximately half in live cells. The major contributing factor to the reduced biotinylation in Δ-Ecad is the increase in *cis*-dimer dissociation when specific *cis* interactions are eliminated. The experimentally determined off-rate (k_off_) for nonspecific *cis* interactions is one order of magnitude higher than the k_off_ for specific *cis* interactions <u>(Thompson et al. 2020)</u>. Consequently, if the Δ mutation significantly reduces the persistence of Ecad *cis* dimers, then there will be less biotinylation, on average, per split-TurboID reconstitution event. Importantly, the Δ-Ecad in the simulations dimerize marginally higher than the values we measure in our experiments since the simulations only capture cumulative binding events and not binding persistence.

The results of our ectodomain binding simulations largely agree with our experimental results, demonstrating that the Δ mutations alone are responsible for the difference in biotinylation. The simulation results of the no-*trans* interaction conditions slightly differ from the experimental results, with the simulations showing a smaller difference in *cis* binding events between WT-Ecad and Δ-Ecad, while the experimental results show a slightly higher incidence of WT *cis* biotinylation compared to Δ cells. While the experimental result can be interpreted to imply that *trans* interactions play a larger role in *cis* binding in Δ cells than in WT cells, the corresponding margins of error are large enough to make such a conclusion unreliable.

In our experiments, we controlled potential sources of binding variation by using A431(EP)-KO cells, where the only classical cadherins present were ones provided by us via transfection. This means that there are no WT-Ecad present at all in Δ cells. Dispase and cell-aggregation experiments with these cells reveal that abolishing specific interactions have significant, detrimental effects on cell-cell adhesion. Our finding runs counter to previous cell aggregation experiments <u>(Harrison et al. 2011),</u> where both WT-Ead and Δ-Ecad expressing cells aggregated to similar extent. A likely reason for this discrepancy is that the A431D cells used in previous cell aggregation experiments were generated using Dexamethasone-induced differentiation instead of targeted CRISPR/Cas9 Ecad/Pcad silencing, and may have still had endogenous populations of WT Ecad and Pcad <u>(Lewis et al. 1997)</u>. Consequently, the presence of WT-cadherins in Δ cells could strengthen cell-cell interactions. Nonetheless, our experiments show that while nonspecific interactions are significantly weaker than specific interactions, the Δ cells still form junctions. This implies a combination of specific and non-specific interactions are responsible for normal Ecad binding behavior and adherens junction formation.

It is important to note that designing a non-*cis* dimerizing negative control is very difficult since it is not possible to fully eliminate Ecad *cis* interactions, because the mechanism underlying nonspecific interactions are not known. Potentially the only reliable way to remove homophilic ectodomain *cis* interactions would be to delete the complete cadherin extracellular region. However, even this drastic approach would not prevent cadherin association through the intracellular domain <u>(Singh et al. 2017)</u>. We therefore designed experiments where we mutated known, specific interactions, to investigate the role of these specific interactions alone while also remaining as close to endogenous conditions as possible. By performing parallel experiments using full-length TurboID, we were able to obtain the proteome for both dimerized and monomeric Ecad, as full-length TurboID biotinylates proximal proteins, independent of their dimerization state. DAT in contrast necessitates that Ecad undergo homophilic *cis* dimerization in order for biotinylation to occur, ensuring that the detected proteome only reflects dimerized Ecad rather than monomeric protein. In this way DAT allows us to separate out dimerization-based interactions from the composite TurboID results.

Our experiments showed that while the core cytoplasmic proteins that associates with Ecad were similar when *cis* dimers were formed via either specific and non-specific contacts, the specific *cis* interactions result in a more robust proteome since Ecads lacking specific interactions were more mobile and form unstable junctions. While WT-DAT returned a tight web of interacting proteins, Δ-DAT showed a broader web of hits, with small unconnected “pockets” of proteins not generally associated with adherens junctions. Furthermore, the adherens junction interaction pathway was significantly downregulated in the Δ cells, indicating less organized junctions and weaker cell adhesion. A protein consistently enriched in WT results over Δ results was Synaptotagmin-like protein 4 (SYTL4) (Fig. S3E, F). This protein is associated with dense core vesicle exocytosis, and is ordinarily thought to inhibit this process <u>(Ostrowski et al. 2010)</u>. The reduction or absence of SYTL4 in the Δ cells could imply an increase in vesicle exocytosis, and further contribute to junctional instability. N-cadherin has been shown to affect dense core vesicle endocytosis in neurons, so it may be that Ecad plays an analogous role in epithelial cells <u>(Dagar et al. 2021)</u>. As no known link exists between Ecad and this process, this could be something that warrants further research. Another interesting result of the STRING guided protein interaction mapping was the presence of a tightly knit group of ribosomal proteins in the DAT results with WT-Ecad, that was not present in the Δ mutant. This could imply that while both WT-Ecad and Δ-Ecad spend time localized to ribosomal units, only WT-Ecad forms dimers in such a way that split TurboID is able to reconstitute. In contrast, lack of specific *cis* interactions prevents this from happening in the Δ mutant.

DAT unfortunately retains TurboID’s lack of total temporal control, as once biotin is added to the cells it is difficult to abort the labeling process in a timely manner. As such, cells must be harvested or stained shortly after the desired labeling period ends, but this is no different from most TurboID experiments. Additionally, if cells cannot be harvested immediately, thorough rinsing with PBS or biotin(-) media can be a way to return cellular biotin concentrations to endogenous levels. The necessity to create cells expressing two populations of modified cadherins could also pose a difficulty, as transfecting and stabilizing such large DNA constructs into the same cells can create heterogeneous expression. However, this can be overcome via multiple rounds of antibody selection, along with correcting for the expression levels of each cadherin during data analysis.

Since DAT requires co-transfection with both EcadC and EcadN, each of which are expressed at different levels, it can be difficult to determine the percentage of total Ecad expressed in a cell that in practice forms a functional DAT pair. For example, EcadC dimerizing with another EcadC would not produce a functional Split TurboID, as each dimer only contains only the C-terminus of Split TurboID. While normalization can be performed by calculating the statistical likelihood of functional dimerization occurring based on relative expression levels, this method does not account for non-specific lateral binding and clustering that creates more complex groups of Ecad bound closely enough that their Split TurboID components can successfully reconstitute. We therefore used a more conservative normalization method by dividing streptavidin intensity by the intensity of the lowest expressing DAT component in the cell line, treating this as a “limiting factor” of potential functional DAT dimerization. This normalized the biotinylation to the maximum amount of functional Split TurboID that could form at one time. In order to compare the two normalization methods, we also analyzed the data by using the statistically determined normalization factor method (Fig. S6A, B) and found that it yielded very similar results to our “limiting factor” method, implying that normalization to the limiting factor can yield robust results.

Other protein dimer-based proximity labeling methods have been developed, such as Split APEX, ContactID, and CATCHFIRE, but these methods both rely on an external agent, e.g. rapamycin, to dimerize the split halves of the labeling enzyme, and also have only been used to investigate *trans* interactions between organelles or cells, and do not have *both* halves of the labeling enzyme on bait proteins <u>(Han et al. 2019; Kwak et al. 2020; Bottone et al. 2023)</u>. Similarly, while we have also previously developed a light-activated TurboID tool, LAB, with high spatial and temporal accuracy and control, here we sought to tie TurboID activation directly to the dimerization state of the bait protein, while LAB cannot differentiate between monomeric and dimeric Ecad <u>(Shafraz et al. 2023)</u>. Another Split-BirA construct, SplitAirID, has been proposed to be useable with two different bait proteins, but has not been experimentally validated <u>(Schaack et al. 2023)</u>. One tool, tSYID, has been developed that utilizes both Split-TurboID and a split fluorophore to create a system where two interacting bait proteins reconstitute both TurboID and the fluorophore to create biotinylation and fluorescence only at the sites of interaction. However, this tool has only been used in plants, and only for cytoplasmic protein baits <u>(Huang et al. 2025)</u>. Our DAT system is thus the first dimer-dependent labeling system developed for use in mammalian cells with large, transmembrane cadherin proteins. We have shown that our approach can be used to directly measure weak and poorly studied Ecad *cis* binding. Non-dimer dependent fluorescence also means DAT can be used as a measure of construct expression without needing a separate reporter.

Since their initial identification, researchers have struggled to fit cadherin *cis* interactions into a comprehensive model. Being much weaker than *trans* interactions, *cis* bonds cannot be measured using established methods such as through solution-phase assays, and the geometry of their bonds excludes them from AFM-based research <u>(Zhang et al. 2009; Häussinger et al. 2002)</u>. Many studies have attempted to characterize these weak yet ubiquitous cadherin interactions, with different investigations coming to sometimes conflicting conclusions <u>(Hong et al. 2010; Wu et al. 2011; Thompson et al. 2021)</u>. By developing a PL tool that reports on *cis* dimerization in live cells, we have been able to resolve that cadherins *cis* dimerize via both specific and nonspecific interactions.

Since the Split-TurboID construct was developed to have low reconstitution affinity in the absence of the desired stimulus <u>(Cho et al. 2020)</u>, we used Split TurboID to make sure biotinylation could be tied to Ecad *cis* dimers and not unrelated Split TurboID reconstitution. In creating a PL tool that only becomes functional when the two halves reconstitute upon dual-bait homodimerization, we have made a spatially and temporally controlled tool engineered for directly studying transmembrane protein *cis* interactions.

## Materials and Methods

### Cloning of Plasmid Constructs

EcadC and EcadN were generated using a cmv-Ecad-EGFP plasmid that was a kind gift from Dr. Soichiro Yamada (University of California, Davis). To generate EcadC, EGFP was removed from the plasmid via restriction digestion with HindIII and NotI to remove the stop codon and create the Ecad-DAT vector. EGFP without the stop codon was generated from cmv-Ecad-EGFP via PCR amplification (primers #1 and #2, Table S1). Split-TurboID-C insert was generated from Addgene plasmid #153003 <u>(Cho et al. 2020)</u> via PCR amplification (primers #3 and #4, Table S1). The Ecad-DAT vector and EGFP and Split-TurboID-C inserts were combined via Gibson Assembly into EcadC. EcadN was generated using the Split-TurboID-N Addgene plasmid #153002 <u>(Cho et al. 2020)</u> linearized via PCR amplification with primers #5 and #6 (Table S1) as a vector. Inserts were generated by amplifying mCherry from Addgene plasmid #36991 <u>(Nekrasova et al. 2011)</u> (primers #7 and #8, Table S1) and Ecad from cmv-Ecad-EGFP (primers #9 and #10, Table S1) using PCR. Vector and inserts were combined using Gibson Assembly.

Δ-EcadC (EcadC with V81D and L175D point mutations on Ecad) was generated via Vector Builder. Δ-EcadN was generated by restriction-digesting the mutated Ecad insert from Δ-EcadC with AgeI and BstEII and inserting into a vector made by restriction digesting EcadN with AgeI and BstEII. Vector and insert were combined via Gibson Assembly.

WT-Ecad fused to full-length TurboID (WT-Turbo) construct was from <u>(Shafraz et al. 2020)</u>. Δ-Turbo was generated by restriction digesting mutated Ecad from Δ-EcadC using enzymes AgeI and BstEII to make a partial Δ-Ecad insert, and using the same enzymes on WT-Turbo to create the vector. Vector and insert were combined via Gibson Assembly.

All primers are listed in Supplementary Table 1.

### Cell Culture and transfection

A431(EP)KO were a kind gift from Dr. Sergey Troyanovsky (Northwestern University). The cells were grown in high-glucose (4.5g/l) Dulbecco’s modified Eagle’s medium (DMEM, Gibco) cell culture medium with 10% fetal bovine serum (Gibco) and 1% penicillin-streptomycin (PSK) (10,000 U/ml, Life Technologies). To generate stable WT-DAT and Δ-DAT cell lines, EcadN and EcadC or Δ-EcadN and Δ-EcadC plasmids were co-transfected into A431(EP)KO using Lipofectamine 3000 (Invitrogen) with equal DNA ratios in a 24-well plate. Cells were passaged after 48 hours and seeded sparsely onto 150mm dishes with 500 μg/mL Geneticin (G418, Gibco) and 5 μg/mL Blasticidin (BSD, Sigma-Aldrich) added to the culture medium for antibiotic selection. Single colonies (∼2mm) were fluorescently screened for dual expression, and those with the brightest and most uniform expression were selected and expanded. To generate the WT-Turbo and Δ-Turbo cell lines, the corresponding plasmids were transfected into A431(EP)KO cells using the same protocol as above and selected with 500 μg/mL Geneticin alone to generate resistant colonies. Colonies were screened for GFP expression, with the brightest and most uniform colony chosen for expansion.

### Immunofluorescence

Cells were grown to confluency and supplemented with 100 µM exogenous biotin for 4 hours for DAT cells. Cells were fixed using 3% paraformaldehyde and 0.3% Triton X-100 in phosphate buffered saline (PBS) for 10 mins and blocked with 1% bovine serum albumin (BSA, VWR Life Science) and 0.3% Triton X-100 in PBS for 30 mins. Anti-GFP antibody (Chicken) and Alexa Fluor 488-conjugated goat anti-chicken-IgY antibody (Invitrogen) were used for GFP-tagged proteins. Anti-mCherry (Rabbit monoclonal, Invitrogen) and Alexa Fluor 568-conjugated goat anti-rabbit-IgG (Invitrogen) were used for mCherry tagged proteins. Alexa Fluor 647-conjugated streptavidin (Invitrogen) was used to label biotin. Primary antibodies were incubated for 1 h (1:1000 dilution in PBS with 1% BSA and 0.3% Triton X-100) and secondary antibodies were incubated for 30 min (1:1000 dilution in PBS with 1% BSA and 0.3% Triton X-100). Cells were imaged using Leica Microsystems Stellaris 5 confocal setup with 63×/1.40 NA oil objective. Images were analyzed using ImageJ.

### Immunofluorescence Image Analysis

Images were analyzed as in <u>(Shafraz et al. 2023)</u> using ImageJ. Briefly, .lif files from the Stellaris 5 were imported as three-channel images for analysis. For confluent cells, ROI’s were chosen by isolating cell membranes expressing both EcadC and EcadN, or Δ-EcadC and Δ-EcadN by creating a threshold mask for the mCherry and GFP detection channels and displaying the overlap between the two channels and then hand-picking individual cell membranes. Intensity values were generated for the mCherry, GFP, and streptavidin channels using the Multi-Measure command. For sparse cell analysis, ROIs were generated by creating masks for the mCherry and GFP channels using the Detect Edges ImageJ functionality, as membrane edges were less clear, and displaying the overlap between the two masks. ROIs were then handpicked, and fluorescence intensity was measured using the Multi-Measure command. For cells exposed to latrunculin, ROIs were generated and measured as with confluent cells with the addition of ROI Area added to the Multi-Measure data collected.

### Diffusion-Reaction Simulations

The initial configuration was generated by randomly distributing molecules on two flat surfaces representing opposed plasma membranes set 24 nm apart <u>(Wu et al. 2010; Boggon et al. 2002)</u>. The system then evolved following a previously developed diffusion-reaction algorithm <u>(Su et al. 2021; Su et al. 2020)</u>. More specifically, a two-step process was used to guide the simulation within each time step. Diffusion was carried out in the first step, whereas the reactions between different types of interactions were implemented sequentially in the second. A Brownian dynamics algorithm was first used to drive Ecad diffusion in each time step by calculating the force between rigid bodies to update their Cartesian coordinates and velocities. Two consecutive rigid bodies of each Ecad were linked with a spring constant to allow conformational variations. The diffusion of the bottom rigid body of each Ecad was confined in its corresponding surface. Finally, a periodic boundary condition along the surface was imposed. The diffusion step was followed by a second step of reaction simulation in which the system checked if there were any new intermolecular bonds formed or if there were any complexes formed between Ecads. Upon Ecad association, a new intermolecular bond was generated to connect the corresponding pair of binding motifs to constrain their diffusion during Brownian dynamics. Upon dissociation, the bond that connected the original complex was broken and the Ecads diffused separately until they encountered another Ecad.

The probabilities of association and dissociation within each simulation step were determined by the reaction rate constants. All association rates in the simulation were fixed to a value that kept the time scale computationally tractable, corresponding to an experimentally measurable *k_on_* of approximately 10^9^ M^-1^s^-1^ which is near the upper limit of diffusion-controlled reactions. The dissociation rate *k_off_* for specific *cis* interactions was set one order of magnitude higher than that for *trans* interactions, while the nonspecific *cis* interactions were assigned a dissociation rate one order of magnitude higher than the specific *cis* interactions. These parameter choices were consistent with previous experimental observations <u>(Thompson et al. 2020)</u>. When both diffusion and reaction steps were completed within a corresponding time step, the new configuration was updated. Finally, as the above process iterated, the kinetics of the system evolved in both Cartesian and compositional space.

### Sample Preparation for MS

A431(EP)KO cells stably expressing WT-DAT, Δ-DAT, WT-Turbo and Δ-Turbo were grown to confluency on 150 mm dishes and supplemented with 100 µM exogenous biotin for 4 hours for DAT cells and 1 hour for Turbo cells. After incubation, cells were washed three times with PBS, scrapped, and centrifuged (775 g for 15 min). Pelleted cells were resuspended in M2 lysis buffer (50 mM Tris-HCl pH 7.5, 150 mM NaCl, 1% SDS and 1% Triton X-100; <u>Muinao et al., 2018</u>) with 5 μl/ml of protease inhibitor mixture (Sigma-Aldrich) and 1 μl/ml benzonase nuclease (250 U/μl; Millipore-Sigma). Cell lysates were flash-frozen in dry ice, rapidly thawed at 37°C, and incubated for 30 min at 4°C. Samples were sonicated at 30% duty cycle for 1 min and centrifuged (14,549 g) for 30 min at 4°C. The supernatants were transferred into a fresh tube, and protein concentration was measured using an RC DC protein assay kit (Bio-Rad). Supernatants (1 mL at 2mg/mL) were incubated with 100 μl super magnetic streptavidin-conjugated beads (Dynabeads MyOne Streptavidin C1, Invitrogen) and rotated overnight at 4°C. The next day, the beads and supernatant were separated. The beads were washed once with lysis buffer, once with wash buffer (2 % SDS in 50mM Tris-HCl at pH 7.4), twice again with lysis buffer, washed three times with 50mM ammonium bicarbonate (NH_4_HCO_3_), and resuspended in 100μl 50mM ammonium bicarbonate containing 4 μg of trypsin for overnight digestion at 37°C in a shaker. The next day, beads were pelleted and separated from the sample using a magnetic holder (Cell Signaling Technology). The remaining tryptic peptides in the supernatant were dried in a vacuum centrifuge. Peptides were analyzed by nanoscale liquid chromatography coupled with tandem mass spectrometry.

### Mass Spectrometry

MS sample analysis was performed as in <u>(Shafraz et al. 2023).</u> Briefly, for each sample, 500ng total peptide was loaded onto a disposable Evotip C18 trap column (Evosep Biosytems, Denmark) and subjected to nanoLC on a Evosep One instrument (Evosep Biosystems). Tips were eluted directly onto a PepSep analytical column, dimensions: 150umx25cm C18 column (PepSep, Denmark) with 1.5 μm particle size (100 Å pores) (Bruker Daltronics). Mobile phases A and B were water with 0.1% formic acid (v/v) and 80/20/0.1% ACN/water/formic acid (v/v/vol), respectively. The standard pre-set method of 60 samples-per-day was used (21minutes chromatography). The MS was done on a hybrid trapped ion mobility spectrometry-quadrupole time of flight mass spectrometer (timsTOF HT, (Bruker Daltonics, Bremen, Germany), operated in PASEF mode. The acquisition scheme used for DIA consisted of 36 precursor windows at width of 25m/z over the mass range 300-1200 Dalton. The TIMS scans layer the doubly and triply charged peptides.

LCMS files were processed with Spectronaut v20 (Biognosys, Zurich, Switzerland) using DirectDIA analysis mode. Mass tolerance/accuracy for precursor and fragment identification was set to default settings. The reviewed FASTA for Homo Sapiens, UP000005640, downloaded from Uniprot on 27/08/2024 and a database of 380 common laboratory contaminants <u>(Frankenfield et al. 2022)</u> were used. Carbamidomethylation of cysteine residues was set as a fixed modification, and methionine oxidation and acetylation of protein N termini as variable modifications. A decoy false discovery rate (FDR) at less than 1% for peptide spectrum matches and protein group identifications was used for spectra filtering (Spectronaut default). Decoy database hits, proteins identified as potential contaminants, and proteins identified exclusively by one site modification were excluded from further analysis.

### Fluorescence Recovery After Photobleaching (FRAP)

WT-Turbo and Δ-Turbo cells were grown to confluency and imaged live. Cells were imaged using Leica Microsystems Stellaris 5 confocal setup with FRAP module enabled and a 63×/1.40 NA oil objective. 1 µm ROIs were placed on the cell junctions at five locations per field of view. Using the FRAP module, bleaching was set for 10 seconds at 1 frame/second with laser power at 200% for each ROI. Imaging proceeded for 250 seconds after bleaching and intensity was recorded once per second. Images were analyzed in ImageJ using the ROIs generated by the microscope FRAP module, with identical control ROIs chosen from the interiors of the cells measured.

### Cell Aggregation Assay

WT-Turbo, Δ-Turbo, and A431(EP)KO cells were used to perform the cell aggregation assay. Cells were seeded at a concentration of 2.5×10^6^ cells/mL. After reaching confluency, cells were washed three times with PBS to remove cell culture media. Cells were then detached using 0.05% Trypsin-EDTA with CaCl₂ supplementation. 1 mM CaCl₂ was added to the trypsin buffer to prevent the digestion of Ecad on the cell surface. After cells were detached, a low-Ca^2+^ suspension buffer (Life Science Ca-free DMEM with 10% Ca-free FBS and 50µm CaCl₂) was used to suspend cells. The cells were centrifuged and rinsed with the suspension buffer. Then 0.1×10^6^ cells/mL were added to each well of a pre-treated 24-well plate with the supplementation of either 2 mM CaCl₂ or 4mM EGTA. The 24-well plate was pre-treated with 0.5% BSA overnight at 4°C to prevent cell attachment to the bottom of the plate. Each well of the plate was washed three times with PBS prior to the start of the experiment. Cells were incubated at 37 °C and 85 rpm for 30 mins. Brightfield images were acquired with an Olympus CK41 microscope.

Analysis was performed using ImageJ. Images were first converted to 8 bit format, and the area of each aggregate was measured using the *Analyze Particles* plugin. Only aggregates that contained at least five or more cells were included in the analysis. The *Analyze Particles* plugin requires a size (pixel^2^) threshold, which was set from 700-infinity. 700 pixels^2^ corresponds on average to an aggregate containing at least five cells.

### Monolayer Dispersion (Dispase) Assay

The monolayer dispersion assay was performed 48 hours after WT-Turbo, Δ-Turbo, and A431(EP)KO cells were seeded at 0.2×10^6^ cells/mL concentration and allowed to reach confluency in a 12-well plate. After three rinses with cell media, cells were incubated with 2.4 U/mL Dispase II (Millipore Sigma) in PBS with MgCl₂ and CaCl₂ for 30 mins at 37 °C. The lifted monolayer was subjected to mechanical agitation for 40 mins at 200 rpm using an orbital shaker. The plate was imaged before and after shaking to count the difference in the number of fragments. Leica Microsystems Stellaris 5 confocal imaging system was used to collect images. Analysis was performed using ImageJ similar to the aggregation assays.

## Supporting information

Reactome Pathways

Signal Comparison Key

## Acknowledgments

Research was supported by the National Institute of General Medical Sciences of the National Institutes of Health (R01GM121885 and R01GM133880) to SS, and (R01GM120238 and R01GM122804) to YW. We thank Dr. Gabriela Grigorean for performing the proteomics sample prep, the LC-MS/MS method writing and running and data analysis in the Proteomics Core Facility of the Genome Center, University of California, Davis. The Bruker timsTOF HT LC/MS system was supported by the Howard Hughes Medical Institute, Investigator Award for Dr. Neal Hunter, UC Davis.

## Author contributions

C.M.O.D. and S.S. designed the study. C.M.O.D., S.S.P., and P.K. performed the experiments and analyzed the data. Y.W. performed and analyzed simulations. S.S. supervised the project. C.M.O.D. and S.S. wrote the manuscript with feedback from all authors.

