## Supplementary Information for "A dimerization-activated proximity labeling system for direct characterization of cadherin *cis* interactions"

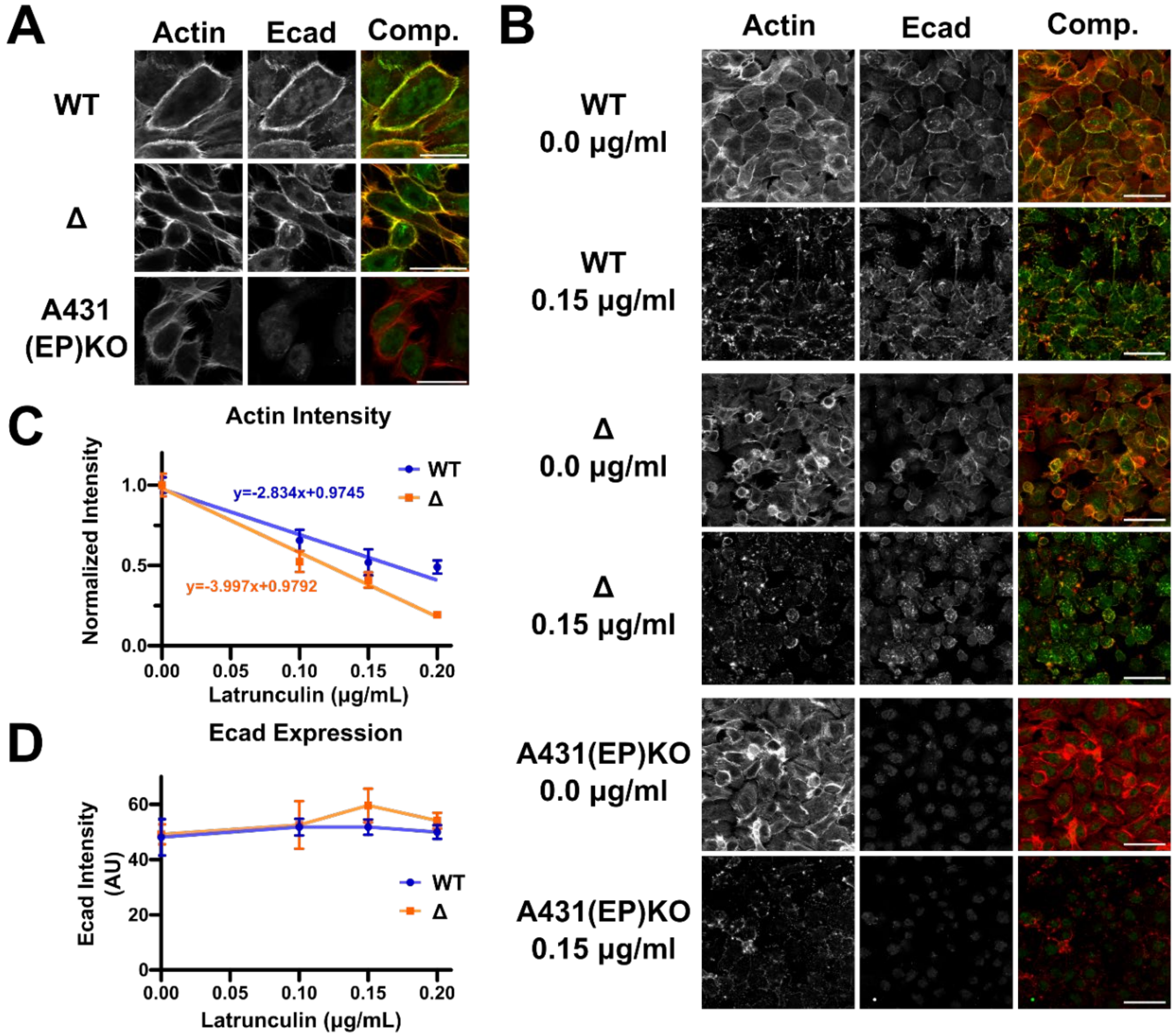


**Supplementary Figure 1: Benchmarking F-actin and its depolymerization. (A)** A431(EP) KO cells expressing WT-Turbo, Δ-Turbo, or only endogenous protein, stained for F-actin using phalloidin, or stained for Ecad. There is strong actin presence along the membrane for both WT-Turbo and Δ-Turbo cells, and weaker actin presence in the parental cells. Scale bars 25 µm. **(B)** A431(EP) KO cells expressing WT-Turbo, Δ-Turbo, or only endogenous protein either without Latrunculin or after incubating in 0.15 µg/ml Latrunculin. Cells are stained for F-actin using phalloidin, or stained for Ecad. Actin staining nearly disappears in the Latrunculin condition. Scale bars 50 µm. **(C)** Actin intensity normalized to Ecad plotted over increasing Latrunculin concentration for WT-Turbo and Δ-Turbo cells. WT-Turbo actin intensity drops to 41.0% of the initial intensity, while Δ-Turbo actin drops to 18.4% of the initial intensity. Measurements taken from n = 8 fields of view (580 $mm^2$) distributed over N = 3 biological replicates. **(D)** Raw Ecad intensity plotted over increasing Latrunculin concentration for WT-Turbo and Δ-Turbo cells. There is no significant change in Ecad density with increasing concentration. Measurements taken from same samples as in (C).

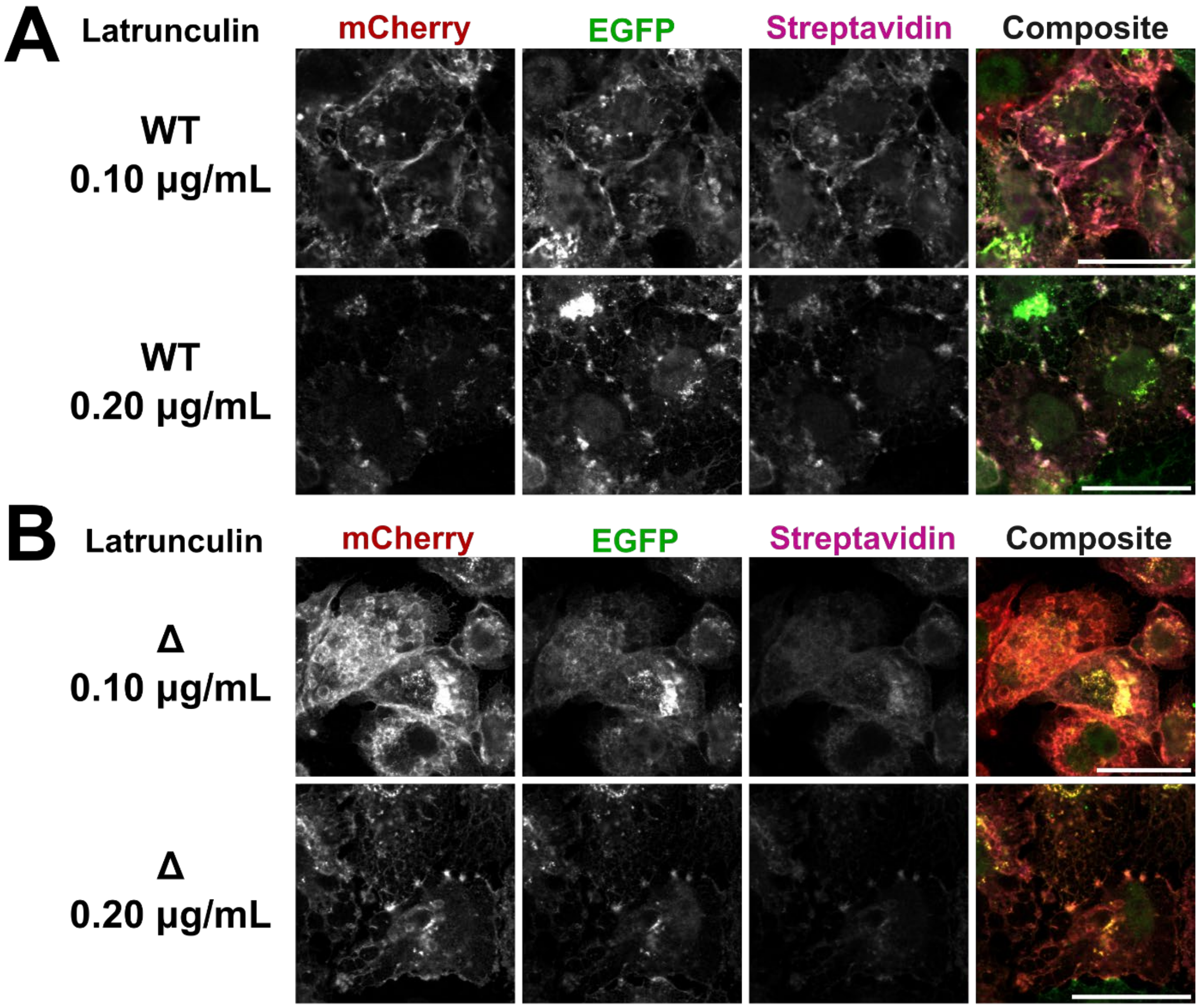


**Supplementary Figure 2: Additional images for Fig. 3C, for Latrunculin concentrations 0.10 and 0.20 µg/ml**. **(A)** WT-DAT and **(B)** Δ-DAT cells. Scale bars are 25 µm.

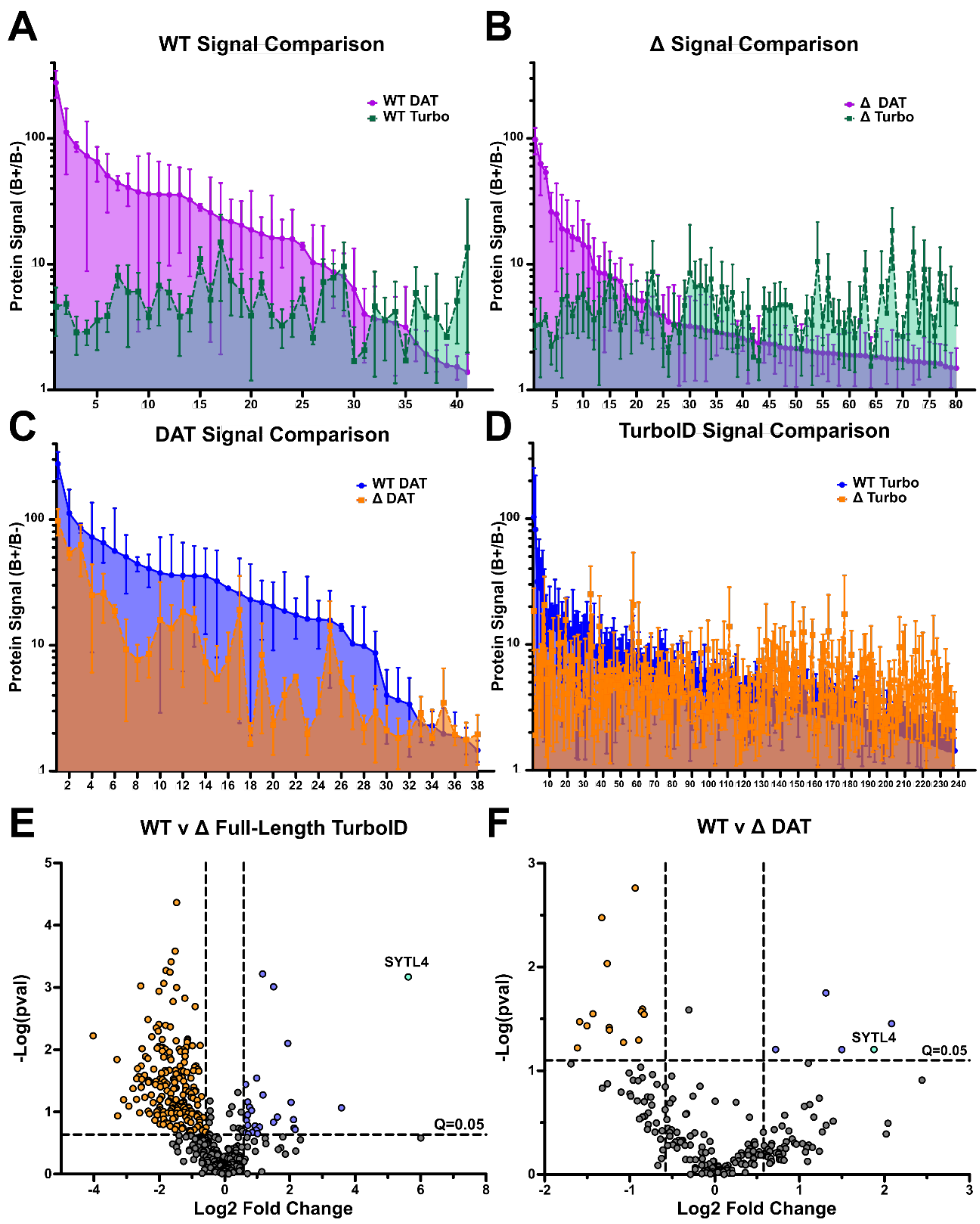


**Supplementary Figure 3: Mass spectrometry signal-to-noise ratios and comparative enrichment of biotinylated proteins. (A-D)** Comparison of mass spectrometry signal-to-noise ratio of proteins shared between different conditions. Comparisons are between WT-DAT and WT-Turbo **(A)**, Δ-DAT and Δ-Turbo **(B)**, WT-Dat and Δ-DAT **(C)**, and WT-Turbo and Δ-Turbo **(D)**. Y-axis values are ratios of protein "quantity" value detected by mass spectrometry for the biotin(+) condition and biotin(-) condition. Results are sorted by WT-DAT values **(A,C)**, Δ-DAT values **(B)**, or WT-Turbo values **(D)**. Protein name keys are in attached Signal Comparison Key document. Error bars are SD. **(E-F)** Volcano plots showing the comparative enrichment of biotinylated proteins in WT-Turbo and Δ-Turbo **(E)** or in WT-DAT and Δ-DAT **(F)**. Proteins with significantly higher enrichment in the WT conditions are colored blue in the upper right quadrant, while proteins with higher enrichment in Δ cells are colored orange in the upper left quadrant. Synaptotagmin-like protein 4 (SYTL4) has been colored cyan.

A) WT-DAT

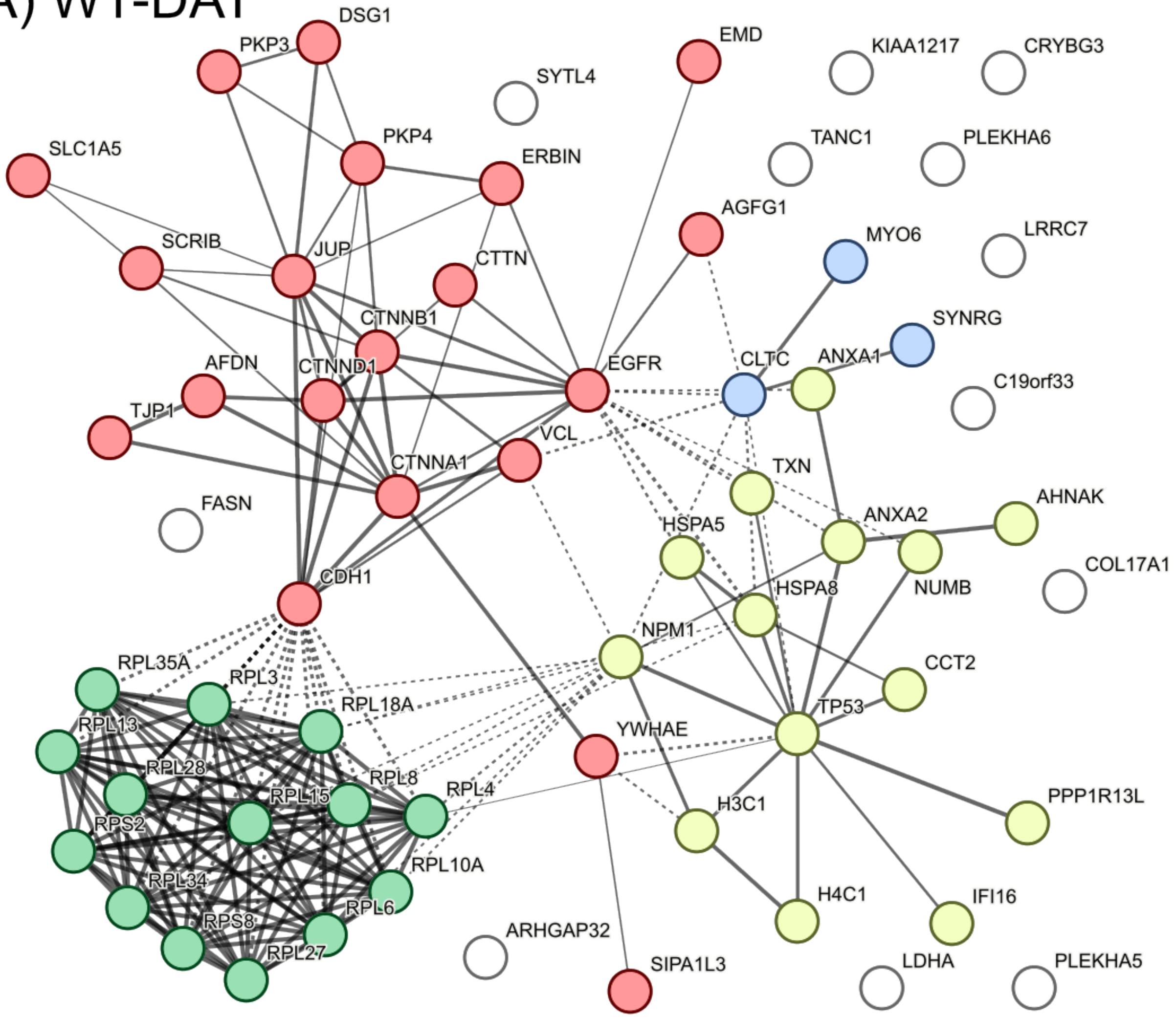


| Color | # Proteins | Group Description |
|---|---|---|
| red | 20 | Cell-cell junction assembly and Adherens junction interactions |
| yellow | 14 | Regulation of gene expression, Cadherin binding |
| green | 14 | Cytoplasmic translation, Ribosome |
| blue | 3 | Gap junction degradation, Clathrin coat of trans-Golgi network vesicle |
| white | 12 | Ungrouped |

**Intra-group Interaction** (solid line)

**Inter-group Interaction** (dashed line)

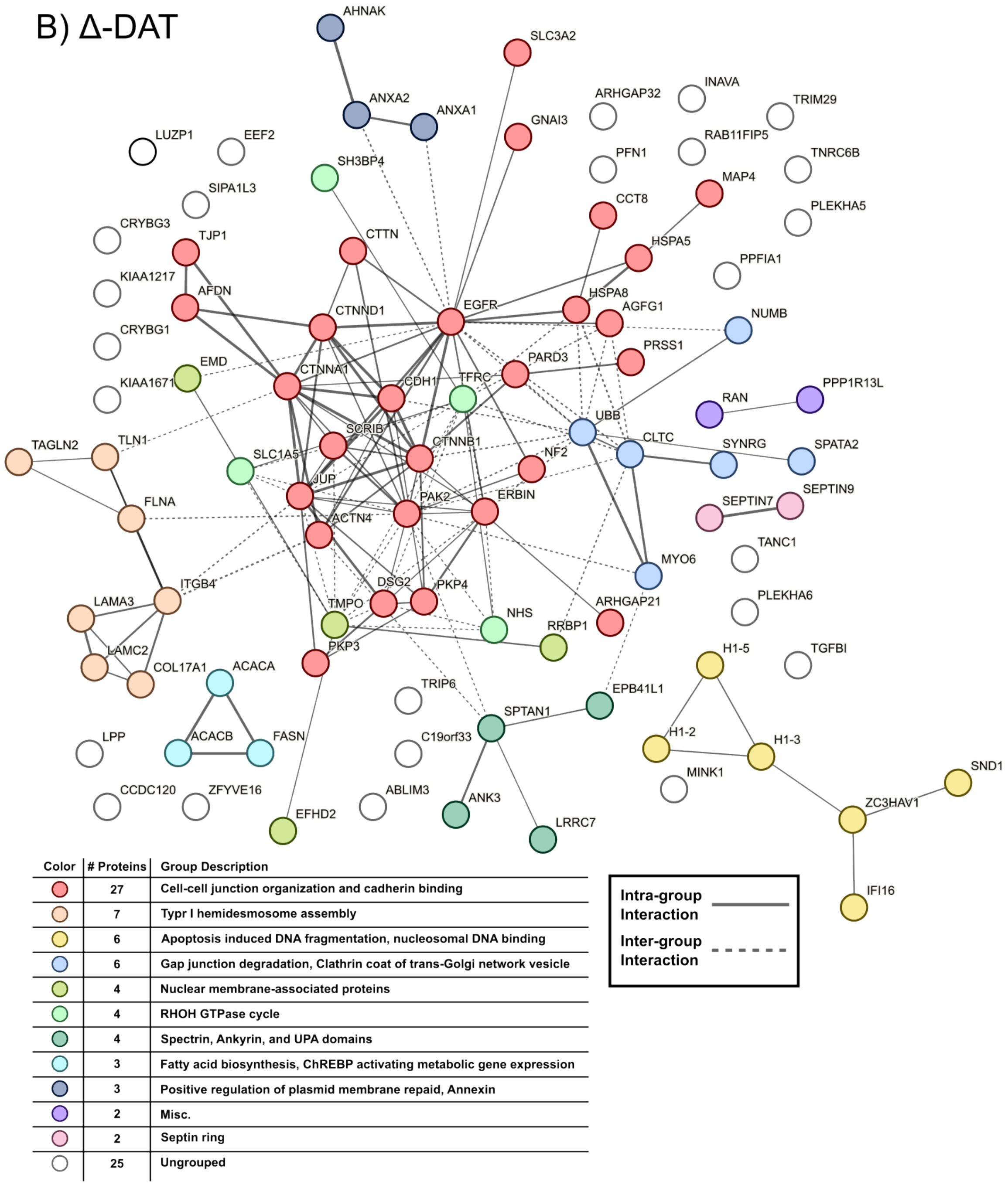


**Supplementary Figure 4: Interaction maps of mass spectrometry hits** for **(A)** WT-DAT and **(B)** Δ-DAT, generated using the STRING online proteomics analysis tool. Line thickness shows confidence of interaction data from Experimental and Database sources. Solid lines represent known interactions between members of the same cluster, while dashed lines represent known interactions between members of different clusters. Clustering was determined using k-means clustering, with the number of clusters set to the maximum number before the software generated single-protein clusters. # clusters = 4 (WT-DAT) or 11 (Δ-DAT). Proteins with no known interactions in the networks are scattered around the outsides of the networks in white.

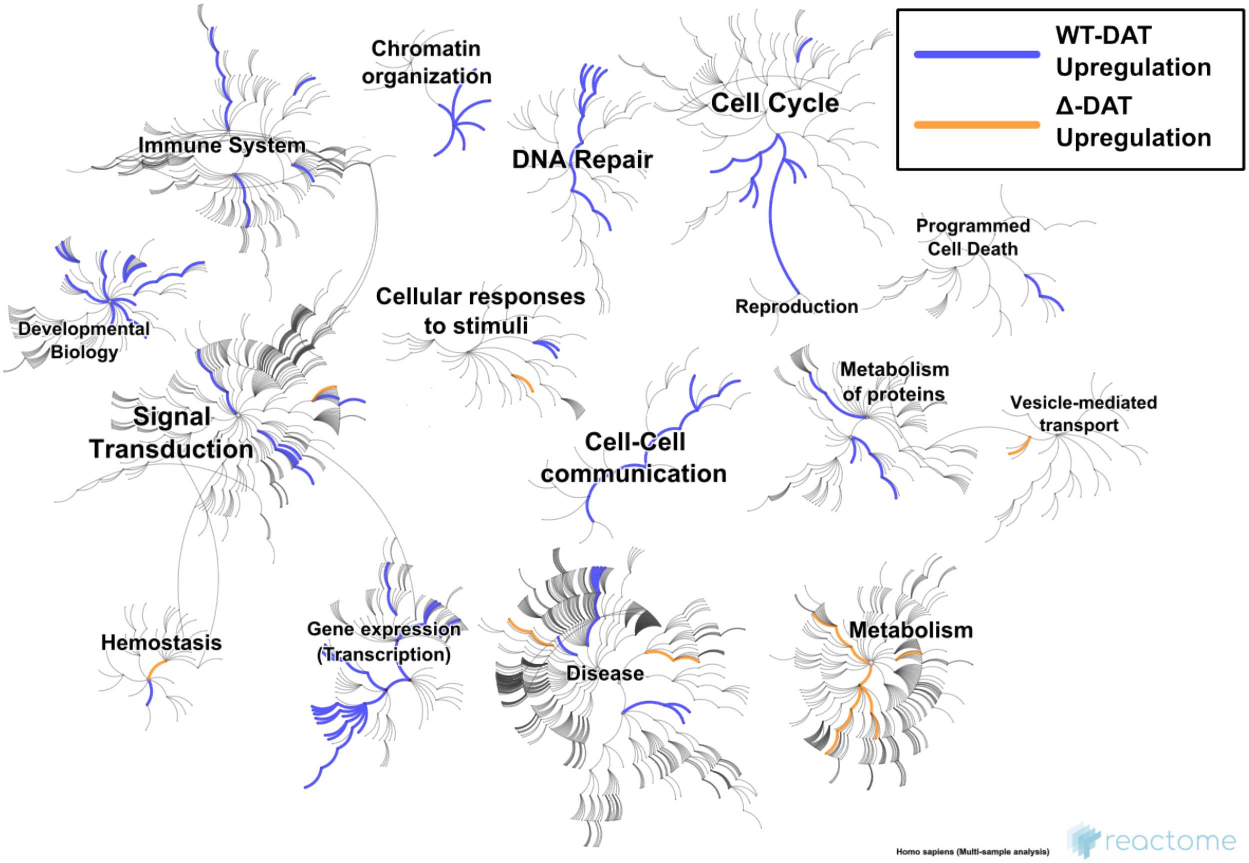


**Supplementary Figure 5: Functional pathway map based on differential protein expression between WT-DAT and Δ-DAT mass spectrometry results, generated in Reactome via the "Analyse gene expression" tool.** The CAMERA analysis method with no additional normalization was used to analyze the proteomics intensity mass spectrometry data. Only pathways with confidence *P < 0.05 are highlighted. Blue pathways were increased in WT-DAT, while orange pathways were increased in Δ-DAT. A comprehensive list of the pathway names is in the Reactome Pathways document.

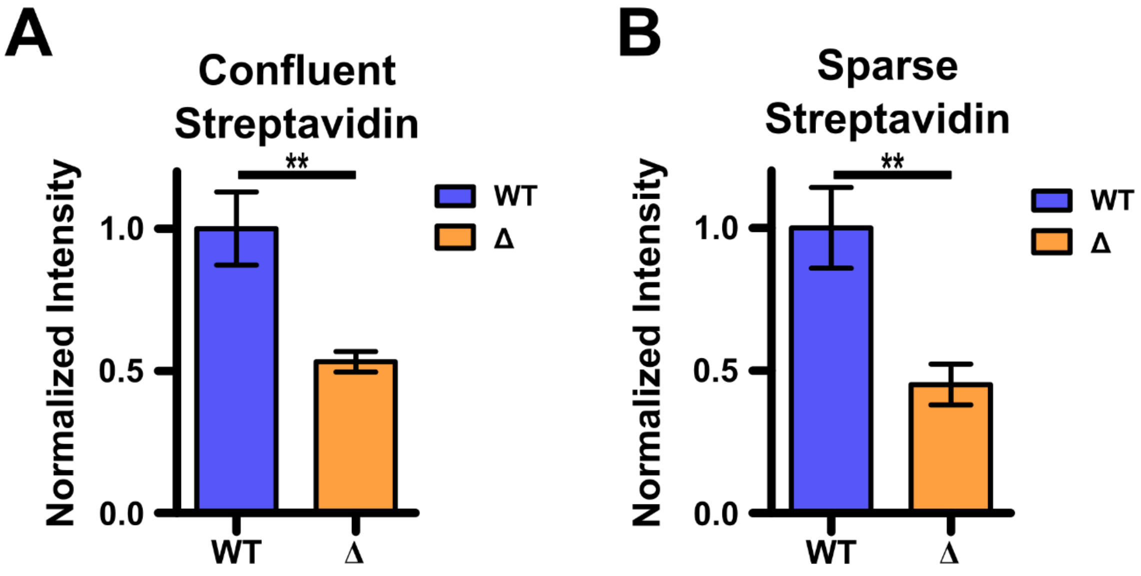


**Supplementary Figure 6: Data analyzed using statistically determined normalization factor method.** Graphs of the data in **(A)** Fig. 1D and **(B)** Fig. 2C analyzed using the statistically determined normalization factor method. The results are comparable to Fig. 1D and Fig. 2C.

**Supplementary Table 1: Primers**

| Primer # | Note | Sequence |
|---|---|---|
| 1 | GFP-F | tggaggtggcgaggacgacctcgagatggtgagcaagggcgaggag |
| 2 | GFP-R | gcccgagcccttaaagcttgacttgtacagctcgtccatg |
| 3 | SptC-F | catggacgagctgtacaagtcaagcttcaagggctcgggctcgacc |
| 4 | SpltC-R | cgcggccgcttagaagctcttttcggcagaccgcagac |
| 5 | EcadN-Vec-F | gctgtacaaggggagtggcagcggaggt |
| 6 | EcadN-Vec-R | gccgtaccgagggcccattttcgaagcctgctttttgtacaa |
| 7 | mCherry-F | tggcgaggacgacctcgagatggtgagcaagggcgagga |
| 8 | mCherry-R | tgccactccccttgtacagctcgtccatgc |
| 9 | Ecad-F | ttgtacaaaaagcaggcttcgaaaatgggccctcggtacggc |
| 10 | Ecad-R | tcctcgcccttgctcaccatctcgaggtcgtcctcgcca |
